# *Salmonella* Typhimurium induces NAIP/NLRC4- and NLRP3/ASC-independent, caspase-1/4-dependent inflammasome activation in human intestinal epithelial cells

**DOI:** 10.1101/2021.12.09.472040

**Authors:** Nawar Naseer, Renate Bauer, Jenna Zhang, Igor E. Brodsky, Isabella Rauch, Sunny Shin

## Abstract

*Salmonella enterica* serovar Typhimurium is a gram-negative pathogen that causes diseases ranging from gastroenteritis to systemic infection and sepsis. *Salmonella* uses type III secretion systems (T3SSs) to inject effectors into host cells. While these effectors are necessary for bacterial invasion and intracellular survival, intracellular delivery of T3SS products also enables detection of *Salmonella* by cytosolic immune sensors. Upon detecting translocated *Salmonella* ligands, these sensors form multimeric complexes called inflammasomes, which activate caspases that lead to proinflammatory cytokine release and pyroptosis. In particular, the *Salmonella* T3SS needle, inner rod, and flagellin proteins activate the NAIP/NLRC4 inflammasome in murine intestinal epithelial cells (IECs), which leads to restriction of bacterial replication and extrusion of infected IECs into the intestinal lumen, thereby preventing systemic dissemination of *Salmonella*. While these processes are studied quite well in mice, the role of the NAIP/NLRC4 inflammasome in human IECs remains unknown. Unexpectedly, we found the NAIP/NLRC4 inflammasome is dispensable for early inflammasome responses to *Salmonella* in both human intestinal epithelial cell lines and organoids. Additionally, the NLRP3 inflammasome and the adaptor protein ASC are not required for inflammasome activation in Caco-2 cells. Instead, we observed a partial requirement for caspase-1, and a necessity for caspase-4 and GSDMD pore-forming activity in mediating inflammasome responses to *Salmonella* in Caco-2 cells. These findings suggest that unlike murine IECs, human IECs do not rely on NAIP/NLRC4, and also do not use NLRP3/ASC. Instead, they primarily use caspases-1 and −4 to mediate early inflammasome responses to SPI-1-expressing *Salmonella*.

## Introduction

Enteric bacterial pathogens such as *Salmonella* enterica serovar Typhimurium (hereafter referred to as *Salmonella*) are leading causes of global morbidity and mortality from diarrheal diseases (1). Contracted upon ingestion of contaminated food or water, *Salmonella* colonizes the intestinal tract, where it uses evolutionarily conserved molecular syringes called type III secretion systems (T3SS) to inject effectors, or virulence factors, into the host cell cytosol (2). *Salmonella* contains two T3SS: the SPI-1 T3SS is expressed early in infection and enables *Salmonella* to invade host cells, while the SPI-2 T3SS is expressed at later timepoints during infection and allows *Salmonella* to replicate within host cells such as intestinal epithelial cells (IECs) (2). IECs thus serve as both the targets of, as well as the first line of physical and innate immune defense against enteric pathogens like *Salmonella*. Most studies of *Salmonella*’s interactions with the innate immune system have been conducted in mice. However, key differences in innate immune genes encoded by mice and humans make it unclear whether mice fully recapitulate how humans respond to *Salmonella*. Here, we interrogated how human IECs sense and respond to *Salmonella* infection.

The mammalian immune system can recognize invading intracellular pathogens through cytosolic sensors such as nucleotide-binding domain, leucine-rich repeat (NLR) receptors. Upon detecting a bacterial ligand or activity, these receptors oligomerize to form multimeric signaling complexes called inflammasomes (3). Inflammasomes recruit and activate cysteine proteases, such as caspase-1 and caspase-8 (3, 4). Some inflammasomes require an adaptor protein called apoptosis-associated speck-like protein containing a CARD (ASC) to mediate their interaction with caspases (3). Active caspases can process the proinflammatory cytokines IL-1α, IL-1β, and IL-18 (3), and the pore-forming protein GSDMD (5). This leads to GSDMD-dependent release of the processed cytokines and an inflammatory form of cell death known as pyroptosis (3). Release of these proinflammatory cytokines alerts nearby cells of the infection, while pyroptosis can eliminate the pathogen’s replicative niche within the infected host cell.

Various cellular insults during infection can trigger activation of different inflammasomes. Inflammasome activation is critical for control of *Salmonella* infection in mice (6). In both murine and human macrophages and murine IECs, *Salmonella* infection activates a family of inflammasome sensors termed NAIPs, which detect the *Salmonella* SPI-1 T3SS needle, inner rod, and flagellin proteins (7–12). Activation of the NAIP/NLRC4 inflammasome specifically in murine IECs restricts bacterial replication, causes extrusion of infected cells from the intestinal epithelial layer, and prevents dissemination of *Salmonella* to systemic sites (13–15). In contrast to mice, humans encode a single NAIP inflammasome sensor, which promiscuously recognizes the T3SS needle, inner rod, and flagellin proteins in human macrophages (10, 16, 17). The role of the NAIP/NLRC4 inflammasome in human IECs during *Salmonella* infection remains unknown.

*Salmonella* infection can also induce the NLRP3 inflammasome, which can be activated by a variety of stimuli during infection, including potassium efflux (18). In murine macrophages, the NLRP3 inflammasome is thought to be important for late timepoints during *Salmonella* infection (19). In murine macrophages and murine and human intestinal epithelial cells, the caspase-11 (mice) or caspase-4/5 (humans) inflammasome is activated late in *Salmonella* infection (19–23). Caspase-4/5/11 detect cytosolic LPS and form the noncanonical inflammasome, which can secondarily activate the NLRP3 inflammasome (24, 25). In human macrophages, *Salmonella* infection triggers recruitment of both NLRC4 and NLRP3 to the same macromolecular complex (26). However, whether the NLRP3 inflammasome plays a functional role during *Salmonella* infection of human IECs has not been previously tested.

Human IECs infected with *Salmonella* undergo caspase-4 inflammasome activation at late time points following infection (10hpi), when there is a considerable population of replicating cytosolic bacteria (20, 22). However, early inflammasome responses to SPI-1-expressing *Salmonella* have not been previously investigated. In this study, we have found that human IECs undergo inflammasome activation in response to SPI-1-expressing *Salmonella*. However, T3SS ligands and flagellin were not sufficient to activate the inflammasome in human Caco-2 cells or intestinal organoids. Additionally, using a combination of pharmacological inhibitors and CRISPR/Cas9 technology, we found that the NAIP inflammasome, the NLRP3 inflammasome, and the adaptor protein ASC are all dispensable for early inflammasome responses to SPI-1-expressing *Salmonella* in Caco-2 cells. Instead, we observed that caspase-1 is partially required, whereas caspase-4 is necessary for inflammasome activation in Caco-2 cells in response to *Salmonella* infection. Our findings delineate the role of several inflammasomes in human IECs *Salmonella* infection. Importantly, these findings indicate how widely inflammasome responses to infection can vary between species as well as cell types.

## Materials and Methods

### Ethics statement

All studies involving human peripheral blood mononuclear cells (PBMCs) and human intestinal organoids were performed in compliance with the requirements of the US Department of Health and Human Services and the principles expressed in the Declaration of Helsinki. Both human PBMCs and organoids are considered to be a secondary use of deidentified human specimens and are exempt via Title 55 Part 46, Subpart A of 46.101 (b) of the Code of Federal Regulations. All experiments performed with murine organoids were done so in compliance with the regulatory standards of, and were approved by the Oregon Health & Science University Institutional Animal Care and Use Committee.

### Bacterial strains and growth conditions

*Salmonella enterica* serovar Typhimurium SL1344 WT and Δ*sipB* (27) isogenic strains were routinely grown shaking overnight at 37°C in Luria-Bertani (LB) broth with streptomycin (100 μg/ml). Cells were infected with *Salmonella* grown under SPI-1-inducing conditions (28).

*Listeria monocytogenes* WT and isogenic strains on the 10403S background were routinely grown shaking overnight at 30°C in brain heart infusion (BHI) broth (29)*. S.* Typhimurium ligands PrgJ or PrgI were translationally fused to the truncated N-terminus of ActA and under the control of the *actA* promoter (29). The *Listeria* strain expressing *S.* Typhimurium SsaI was constructed using a codon-optimized gene fragment (IDT) cloned into the pPL2 vector and introduced into *Listeria* as previously described (29, 30).

### Bacterial infections

Where indicated, cells were primed with 100ng/ml or 400ng/ml of Pam3CSK4 (Invivogen) for 3 hours prior to infection. To induce SPI-1 expression, overnight cultures of *Salmonella* were diluted into LB broth containing 300 mM NaCl and grown for 3 hours standing at 37°C (28). Overnight cultures of *Listeria* were diluted and grown shaking for 3 hours in BHI. All cultures were pelleted at 6,010 × *g* for 3 minutes, washed once with PBS, and resuspended in PBS. Cells were infected with *Salmonella* at a multiplicity of infection (MOI) of 60 or *Listeria* at the indicated MOI for each experiment in the figure legend. Infected cells were centrifuged at 290 × *g* for 10 minutes and incubated at 37°C. 1 hour post-infection, cells were treated with 100ng/mL or 50ng/mL of gentamicin to kill any extracellular *Salmonella* or *Listeria* respectively. Infections proceeded at 37°C for the indicated length of time for each experiment. Control cells were mock-infected with PBS for all experiments.

### Cell culture of intestinal epithelial cell lines

All cell lines were obtained from American Type Culture Collection (ATCC). Caco-2 cells (HTB-37; ATCC) were maintained in DMEM supplemented with 10% (vol/vol) heat-inactivated FBS, 100 IU/mL penicillin, and 100 μg/mL streptomycin. T84 cells (CCL-248; ATCC) were maintained in DMEM F-12 supplemented with 5% (vol/vol) heat-inactivated FBS, 100 IU/mL penicillin, and 100 μg/mL streptomycin.

One day prior to infection or treatment, cells were dissociated with 0.25% Trypsin-EDTA (Gibco) diluted 1:1 with 1X PBS. Cells were incubated with trypsin at 37°C for 15 minutes, after which the trypsin was neutralized with serum-containing media. Cells were replated in media without antibiotics in a 24-well plate at a concentration of 3 × 10^5^ cells/well. Where indicated, cells were primed with 100ng/mL or 400ng/ml Pam3CSK4 (Invivogen) or 500ng/ml LPS (Sigma-Aldrich) for 3h or 16h prior to anthrax toxin treatment or bacterial infections.

### Culture of intestinal organoids in spheroid culture

Spheroids derived from human duodenum or colon were kindly provided by Jared Fisher at Oregon Health & Science University. Human and murine organoids were cultivated in special conditioned medium as described by Myoshi & Stappenbeck (31). Briefly, spheroids were grown in Corning® Matrigel® Basement Membrane Matrix (VWR). Spheroids were dissociated with Trypsin-EDTA (1X PBS, 0.25% trypsin, 0.5 mM EDTA) at 37°C for 5 minutes. Subsequently, washing medium (DMEM/F12 with HEPES (Sigma), 10% FBS (heat-inactivated), 1X P/S) was added to stop trypsinization. After washing the cells with washing medium at 200 × *g* for 5 minutes at room temperature (RT), the supernatant was completely aspirated and the cell pellet resuspended in Matrigel®. 20 µL matrigel-cell drops were placed into 24-well plates and polymerized for 5 minutes at 37°C upside down. Afterwards, cells were cultured in organoid medium consisting of 50% primary culture medium (Advanced DMEM/F12 (Gibco), 20% FBS, 1X P/S, 1X L-Glu) and 50% conditioned medium (L-WRN-cell supernatant) supplemented with 10 µM of ROCK inhibitor Y27632 (Sigma) and 10 µM TGF-β inhibitor SB431542 (Millipore) (for murine organoids only).

### ELISAs

Supernatants harvest from infected cells were assayed using ELISA kits for human IL-18 (R&D Systems) and IL-8 (R&D Systems).

### Immunoblot analysis

Cell lysates were harvested for immunoblot analysis by adding 1X SDS/PAGE sample buffer to cells following infection. Cells were incubated and infected in serum-free media to collect supernatant samples. Supernatant samples were centrifuged at 200 × *g* to pellet any cell debris. The supernatant was then treated with trichloroacetic acid (TCA) (25 µL of TCA for 500 µL of supernatant) overnight at 4°C. The next day, the samples were centrifuged at maximum speed (15871 × *g*) for 15 minutes at 4°C. Precipitated supernatant pellets were washed with ice-cold acetone, centrifuged at maximum speed (15871 × *g*) for 10 minutes at 4°C, and resuspended in 1X SDS/PAGE sample buffer. All protein samples (lysates and supernatants) were boiled for 5 minutes. Samples were separated by SDS/PAGE on a 12% (vol/vol) acrylamide gel, and transferred to PVDF Immobilon-P membranes (Millipore). Primary antibodies specific for human IL-18 (MBL International PM014) and β-actin (4967L; Cell Signaling) and HRP-conjugated secondary antibody anti-rabbit IgG (7074S; Cell Signaling) were used. ECL Western Blotting Substrate or SuperSignal West Femto (both from Pierce Thermo Scientific) were used as the HRP substrate for detection.

### Propidium iodide (PI) uptake assay

7.5 × 10^4^ Caco-2 cells per well were plated in a black, flat-bottom 96-well plate (Cellstar) in PI uptake media containing 1X HBSS without phenol red, 20 mM HEPES, and 10% (vol/vol) heat-inactivated FBS. Cells were infected at an MOI=60 and control wells were treated with 1% Triton. After infection, cells were centrifuged at 290 × *g* for 10 minutes following infection. 5 μM propidium iodide (PI, P3566, Invitrogen) diluted in PI uptake media was added to the cells. The plate was sealed with adhesive optical plate sealing film (Microseal, Bio-Rad) and placed in a Synergy H1 microplate reader (BioTek) pre-heated to 37°C. PI fluorescence was measured every 10 minutes for the indicated number of hours.

### Anthrax toxin-mediated delivery of bacterial ligands into organoids

Organoids were seeded in 96-well plates with transparent bottom and opaque walls in 5µl Matrigel domes. After two days in organoid media, organoids were grown for an additional 3 days in differentiation media: DMEM/F12 supplemented with 20% murine R-Spondin1 supernatant, 10% murine Noggin supernatant, 50 ng/mL recombinant murine EGF (Fisher Scientific), 1X P/S, 1X L-Glu, 10 mM HEPES (HiMedia), 1X N2 (Life Technologies), 1X B27 (Life Technologies), and 1 mM N-acetylcysteine (Fisher Scientific). 5 µM DAPT was added for the last 24 hours.

Differentiated organoids were then treated with 16µg/ml PA and 8µg/ml LFn-FlaA or 0.1 µg/ml LFn-PrgJ in differentiation media containing 10µg/ml propidium iodide for 4 hours. Total lysis wells were treated with 1%Triton.

### Expression of inflammasome genes in human small intestinal organoids

To analyze inflammasome expression under different media conditions, spheroids were cultured in differentiation medium: DMEM/F12 supplemented with 20% supernatant from R-Spondin1 expressing L-cells, 10% supernatant from Noggin expressing cells, 50 ng/mL recombinant murine EGF (Fisher Scientific), 1X P/S, 1X L-Glu, 10 mM HEPES (HiMedia), 1X N2 (Life Technologies), 1X B27 (Life Technologies), 1 mM N-acetylcysteine (Fisher Scientific), and 5 µM DAPT. After 4 days of incubation at 37°C, Matrigel domes were dissolved in PBS-EDTA (5 mM) for 1 hour at 4°C on an orbital shaker. After centrifugation at 300 × *g* for 5 minutes at 4°C, the cell pellet was resuspended in TRIzol^TM^ to analyze mRNA expression.

### Isolation of peripheral blood mononuclear cells (PBMCs)

To compare expression levels of inflammasome components in human intestinal epithelial cells with immune cells, PBMC-derived cDNA was kindly provided by William Messer at Oregon Health & Science University. Briefly, PBMCs were isolated using density gradient centrifugation. After overlay of Lymphoprep^TM^ (Alere Technologies AS) with blood mixed 1:2 with 1X PBS (pH 7.4, Gibco), the sample was centrifuged at 800 × *g* for 20 minutes at room temperature (RT). Residual erythrocytes were lysed with 1X RBC lysis buffer (10X, BioLegend), followed by three washing steps with fresh PBS for 10 minutes at 250 × *g*. Subsequently cells were harvested in TRIzol^TM^ Reagent (Thermo Fisher Scientific) for mRNA analysis.

### RNA extraction, cDNA synthesis, and real-time quantitative polymerase chain reaction (RT-qPCR) of organoid and PBMC samples

After thawing the TRIzol^TM^ samples, chloroform was added and the tubes centrifuged for 15 minutes at 12,000 × *g* at 4°C. The aqueous phase was transferred to a new tube containing linear polyacrylamide (Gene-Elute^TM^ LPA, Sigma). To allow RNA precipitation, the samples were incubated with isopropanol for 10 minutes and subsequently centrifuged for 10 minutes at 12,000 × *g*. After aspiration of the supernatant, the RNA pellet was washed once with 75% ethanol for 5 minutes at 7,500 × *g* at 4°C. The supernatant was aspirated and the dried pellet resuspended in Molecular Biology Grade Water (Corning). After determination of RNA content and quality (260/280 and 260/230 ratios), 1 µg of RNA was reversely transcribed into cDNA. Reaction steps were performed in a Biorad T100 Cycler. First, residual DNA was removed using RQ1 RNAse-free DNase (Promega) in RQ1 DNase 1X Reaction buffer (Promega) for 30 minutes at 37°C. After stopping the reaction with RQ1 DNase Stop Solution (Promega) for 10 minutes at 65°C, Oligo dTs (Sigma) and dNTPs (Sigma) were added for 5 minutes at 65°C. Reverse transcription was performed with SuperScript^TM^ IV Reverse Transcriptase (Invitrogen), 5 mM DTT and SuperScript^TM^ IV Reaction Buffer (Invitrogen) for 10 minutes at 50-55°C. Subsequently the enzyme was inactivated at 80°C for 10 minutes.

To analyze RNA expression, cDNA was mixed with 10 µM forward and reverse primers and PowerUp^TM^ SYBR^TM^ Green Master Mix (Applied Biosystems) according to manufacturer’s instructions. The following primers were used:

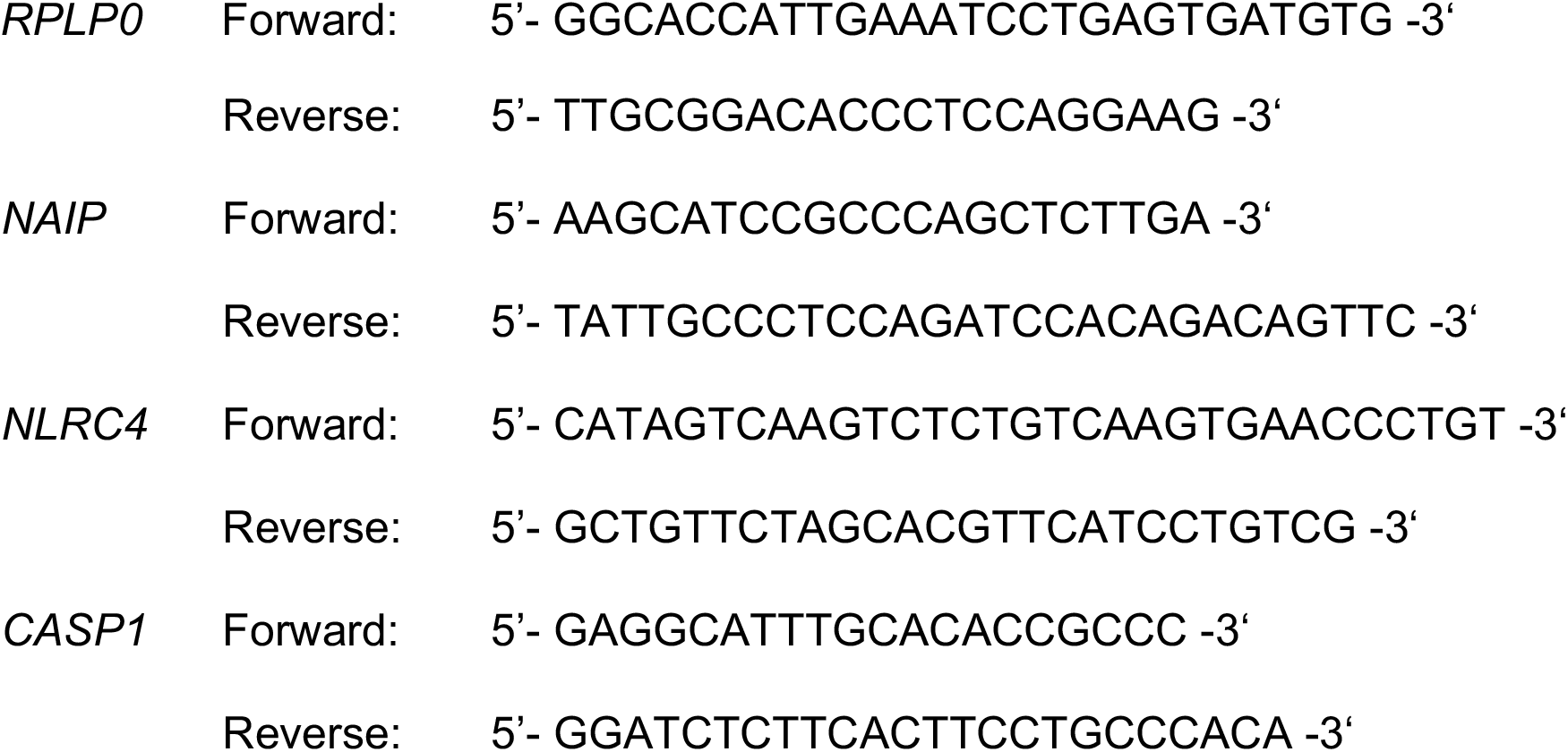

Amplification was analyzed in real-time with StepOne Software v2.3. In brief, samples were incubated for 10 minutes at 95°C, followed by 40 cycles of heating to 95°C for 15 seconds and cooling to 60°C for 1 minute. To monitor specificity of the run, the melt curves were determined by keeping the samples at 95°C for 15 seconds, cooling to 60°C for 1 min and then increasing the temperature every 15 seconds by 0.3°C up to 95°C. Expression levels relative to the housekeeping gene (*RPLP0*) were calculated using the formula x = 2^-Δct^.

### RNA extraction, cDNA synthesis, and RT-qPCR of Caco-2, T84, and THP-1 samples

RNA was isolated using the RNeasy Plus Mini Kit (Qiagen) following the manufacturer’s instructions. Cells were lysed in 350 μL RLT buffer with β-mercaptoethanol and centrifuged through a QIAshredder spin column (Qiagen). cDNA was synthesized from isolated RNA using SuperScript II Reverse Transcriptase (Invitrogen) following the manufacturer’s protocol. Quantitative PCR was conducted with the CFX96 real-time system from Bio-Rad using the SsoFast EvaGreen Supermix with Low ROX (Bio-Rad). To calculate relative gene expression, mRNA levels of target genes were normalized to housekeeping gene *HPRT* and the formula x = 2^−ΔCT^ was used. The following primers from PrimerBank (PrimerBank identification listed within parentheses) were used (32–34):

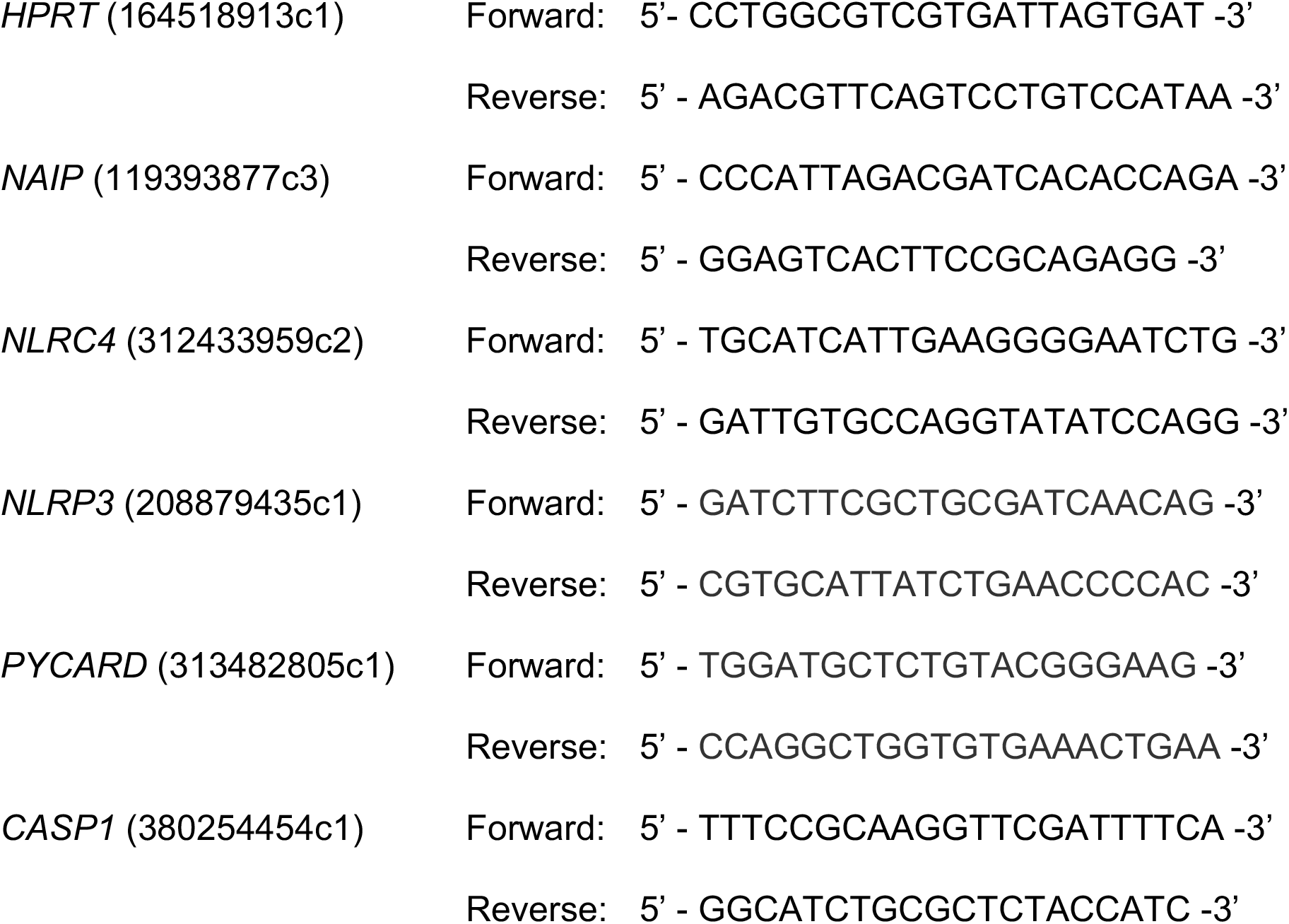

### Inhibitor experiments

Cells were treated 1 hour prior to infection at the indicated concentrations with inhibitors: varying concentrations of MCC950 (Sigma Aldrich PZ0280), 20 μM of pan-caspase inhibitor Z-VAD(OMe)-FMK (SM Biochemicals SMFMK001), 25 μM of caspase-1 inhibitor Ac-YVAD-cmk (Sigma Aldrich SML0429), 30 μM of disulfiram (Sigma).

### siRNA-mediated knockdown of genes

The following Silencer Select siRNA oligos were purchased from Ambion (Life Technologies): *CASP4* (s2412), *CASP5* (s2417), two Silencer Select negative control siRNAs (Silencer Select Negative Control No. 1 siRNA and Silencer Select Negative Control No. 2 siRNA). Three days before infection, 30 nM of siRNA was transfected into Caco-2s using Lipofectamine RNAiMAX transfection reagent (Thermo Fisher Scientific) following the manufacturer’s protocol. Cells were primed with 400 ng/ml of Pam3CSK4 (Invivogen) for 3 hours prior to infection.

## Statistical analysis

Prism 9.1.1 (GraphPad Software) was used to graph all data and for all statistical analyses. Statistical significance for experiments was determined using the appropriate test and are indicated in each figure legend. Differences were considered statistically significant if the *p* value was <0.05.

## Results

### *Salmonella* infection induces inflammasome activation in human intestinal epithelial cells

Once inside the host, *Salmonella* upregulates expression of its SPI-1 T3SS, which delivers effectors that enable *Salmonella* to invade intestinal epithelial cells (2). Human IECs infected with *Salmonella* undergo inflammasome activation at late time points following infection (10hpi) (20, 22). However, early inflammasome responses to SPI-1-expressing *Salmonella* have not been previously investigated. To test if human IECs undergo early inflammasome activation in response to *Salmonella* grown under SPI-1-inducing conditions (28), we infected WT Caco-2 cells, a human colorectal cell line, with WT *Salmonella* (WT Stm) or *Salmonella* lacking its SPI-1 T3SS (Δ*sipB* Stm) and assayed for subsequent inflammasome activation by measuring release and cleavage of the inflammasome-dependent cytokine IL-18 at 6hpi, an early time point following *Salmonella* infection (Fig. 1A, B). Cells infected with WT Stm released significantly increased levels of cleaved IL-18 into the supernatant compared to mock-infected cells (Fig. 1A, B). In contrast, cells infected with Δ*sipB* Stm, which is unable to invade cells, failed to release cleaved IL-18 (Fig. 1A, B). We measured cell death as another readout of inflammasome activation by assaying for uptake of the cell-impermeable dye propidium iodide (PI), which enters cells as they form pores in their plasma membrane and undergo cell death. In cells infected with WT Stm, PI uptake began to occur between 4-6hpi, and gradually increased over time, indicating that infected cells begin to undergo cell death as early as 4-6hpi. As expected, cells infected with Δ*sipB* Stm, which cannot enter the host cell, did not uptake any PI. We found that inflammasome activation in response to *Salmonella* also occurs in another human colorectal cell line, T84 cells (Fig. 1D). T84 cells infected with SPI-1-expressing WT Stm also released IL-18 into the supernatant at 6hpi (Fig. 1D). As a control, we found that in both Caco-2 and T84 cells, the inflammasome-independent cytokine, IL-8, is released in response to WT and Δ*sipB* Stm (Fig. S1A, S1B).

**Figure 1:**
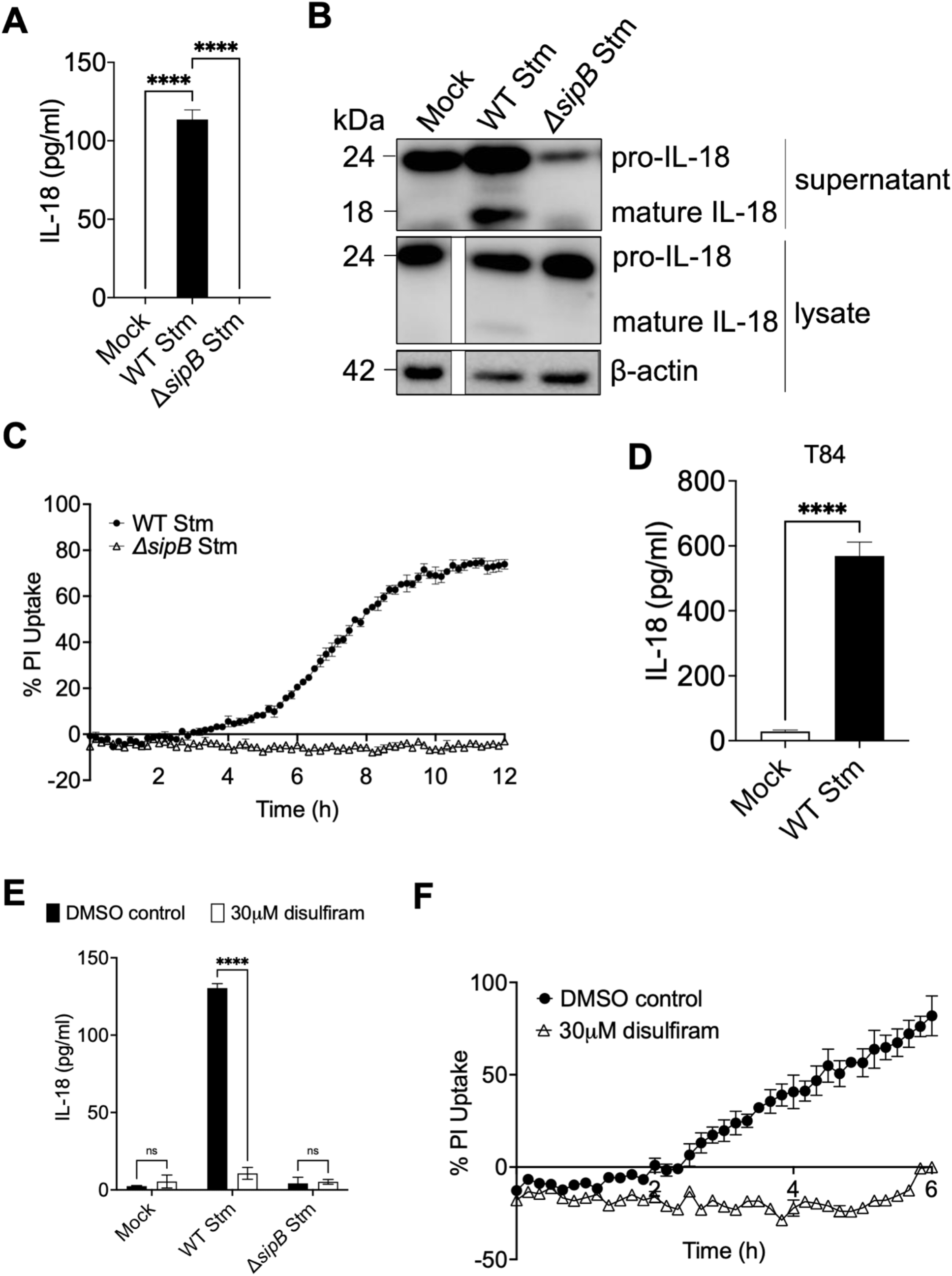
*Salmonella* infection induces inflammasome activation in human intestinal epithelial cells. Caco-2 cells (A – C, E, F) or T84 cells (D) were infected with PBS (Mock), WT *S*. Typhimurium, or Δ*sipB S*. Typhimurium. (A, D) Release of IL-18 into the supernatant was measured by ELISA at 6hpi. (B) Lysates and supernatants collected 6hpi were immunoblotted for IL-18 and β-actin. (C) Cell death was measured as percentage uptake of propidium iodide, normalized to cells treated with 1% Triton. (E, F) Caco-2 cells were treated with 30 μM disulfiram or DMSO as a vehicle control 1 hour prior to infection. Cells were then infected with PBS (Mock), WT *S.* Typhimurium, or Δ*sipB S*. Typhimurium. (E) Release of IL-18 into the supernatant was measured by ELISA at 6hpi. (F) Cell death as percentage uptake of propidium iodide, normalized to cells treated with 1% Triton. ns – not significant, **** *p* < 0.0001 by Dunnett’s multiple comparisons test (A), or by unpaired t-test (C, D, F) or by Šídák’s multiple comparisons test (E). Error bars represent the standard deviation or the standard error of the mean (SEM) (C, F) of triplicate wells from one experiment. Data shown are representative of at least three independent experiments.

Inflammasome activation leads to cleavage of the pore-forming protein GSDMD (5, 35). The N-terminal fragment of cleaved GSDMD inserts into the host cell plasma membrane and oligomerizes to create a pore through which cellular components such as cleaved IL-1 and IL-18 can be released (36–39). These pores eventually cause rupture of the cell through osmotic flux, resulting in pyroptosis (3). To determine if pore formation by GSDMD is required for release of IL-18 and PI uptake in human IECs infected with *Salmonella*, we pretreated Caco-2 cells with disulfiram, which prevents cleaved GSDMD from inserting into the plasma membrane and thus abrogates pore formation (40). Treatment with disulfiram led to loss of IL-18 release and PI uptake during *Salmonella* infection (Fig. 1E, 1F). Importantly, release of the inflammasome-independent cytokine IL-8 was not affected by disulfiram treatment (Fig. S1C).

Collectively, these results suggest that as early as 4-6hpi, human IECs infected with WT Stm grown under SPI-1-inducing conditions undergo inflammasome activation, and GSDMD-mediated pore formation is required to observe IL-18 release and PI uptake in response to *Salmonella* in human IECs.

### Bacterial T3SS ligands do not activate the inflammasome in human intestinal epithelial cells

We next sought to determine the bacterial ligands that trigger inflammasome activation in human IECs. The *Salmonella* SPI-1 T3SS inner rod protein (PrgJ), the SPI-1 T3SS needle protein (PrgI), and flagellin activate the NAIP/NLRC4 inflammasome in both murine and human macrophages, (8, 11). Given that *Salmonella* grown under SPI-1-inducing conditions triggered inflammasome activation in human IECs (Fig. 1), we hypothesized that this inflammasome activation was due to NAIP-mediated recognition of *Salmonella* T3SS ligands. First, we asked whether *Salmonella* T3SS ligands are sufficient to activate the inflammasome in human IECs. We used the Gram-positive bacterium *Listeria monocytogenes* to deliver *Salmonella* T3SS ligands into IECs, We have previously used the *Listeria* system to deliver *Salmonella* T3SS ligands into the cytosol of human macrophages to examine NAIP-dependent inflammasome activation (17). In this system, ligands of interest are translationally fused to the N-terminus of truncated ActA, enabling the ligands to be delivered into the host cell cytosol (17, 29).

Human macrophages infected with *Listeria* expressing the *Salmonella* SPI-1 T3SS inner rod PrgJ or needle PrgI undergo robust inflammasome activation (17). Surprisingly, IECs infected with *Listeria* expressing either the SPI-1 inner rod (PrgJ) or needle (PrgI) proteins failed to induce IL-18 release (Fig. 2A, B; Fig. S2A, B). Polarized Caco-2 cells and C2Bbe1 cells, a Caco-2 subtype, infected with *Listeria* expressing the SPI-1 inner rod PrgJ also failed to release IL-18 levels above that observed in cells infected with WT Lm (Fig. S2C, D). Collectively, these data indicate that *Listeria* delivery of bacterial T3SS ligands is not sufficient to induce inflammasome activation in human IEC lines.

**Figure 2:**
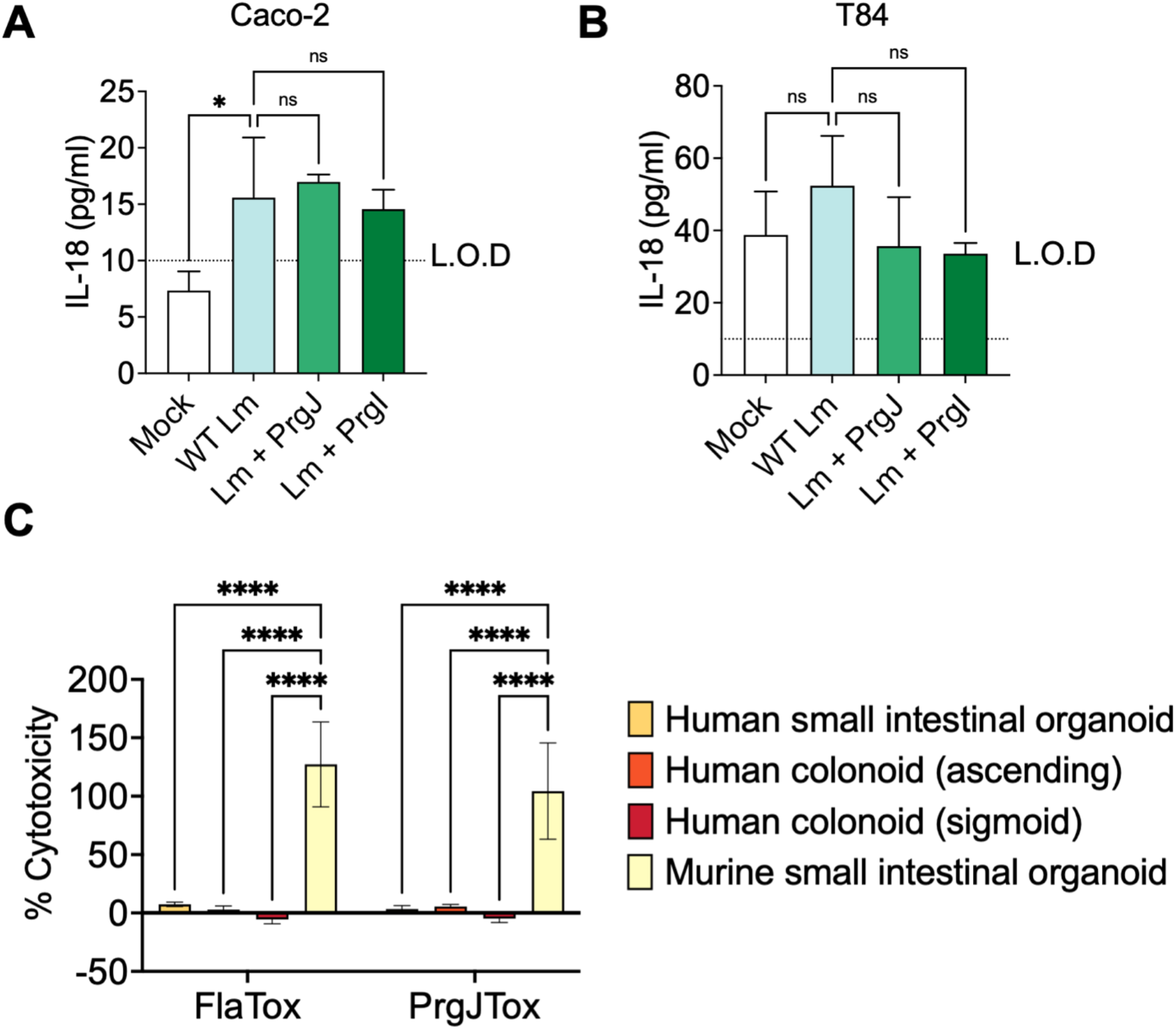
Bacterial T3SS ligands do not activate the inflammasome in human intestinal epithelial cells. (A, B) Caco-2 (A) or T84 cells (B) were primed for 3h with 100 ng/ml of Pam3CSK4 and infected with PBS (Mock), WT *L. monocytogenes* (WT Lm), or *L. monocytogenes* expressing *S.* Typhimurium SPI-1 inner rod (Lm + PrgJ), or SPI-1 needle (Lm + PrgI), at an MOI of 100. Release of IL-18 into the supernatant was measured by ELISA at 16hpi. L.O.D indicates the limit of detection of the assay. (C) Differentiated intestinal organoids were treated with FlaTox (PA + LFn-Fla) or Inner Rod Tox (PA + LFn-Rod) in media containing propidium iodide for 4h. Cell death was measured as percentage uptake of propidium iodide, normalized to organoids treated with 1% Triton. ns – not significant, * *p* < 0.05, **** *p* < 0.0001 by Dunnett’s multiple comparisons test. Error bars represent the standard deviation of triplicate wells from one experiment. Data shown are representative of at least three independent experiments.

As we did not observe inflammasome activation by bacterial T3SS ligands delivered using *Listeria*, we tested a second delivery method, the *Bacillus anthracis* toxin system, to deliver these bacterial ligands into the cytosol of IECs (41). The anthrax toxin delivery system contains two subunits: a protective antigen (PA) that creates a pore in the host endosomal membrane and a truncated lethal factor (LFn) that is delivered through the PA pore into the cytosol. T3SS ligands that are translationally fused to the N-terminal domain of the *B. anthracis* LFn are delivered into the host cell cytosol upon treatment with both PA and the LFn fusion (41) (collectively referred to as Tox). We delivered the *Salmonella* SPI-1 T3SS inner rod protein (PrgJTox) into Caco-2 cells (Fig. S3A), polarized Caco-2 cells (Fig. S3B), C2Bbe1 cells (Fig. S3C), and T84 cells (Fig. S3D) and assayed the release of the inflammasome-dependent cytokine IL-18. In all cell types, we failed to observe IL-18 secretion in response to PrgJTox that was above the PA alone control (Fig. S3). Thus, our data indicate that human IEC lines did not undergo inflammasome activation in response to bacterial T3SS ligands delivered with the anthrax toxin system.

To determine if this absence of inflammasome responses to T3SS ligands was limited to immortalized IECs or extended to non-immortalized IECs as well, we delivered flagellin (FlaTox) or the SPI-1 T3SS inner rod PrgJ (PrgJTox) into human intestinal organoids and measured levels of cell death (Fig. 2C). Both human small intestinal organoids and colonoids failed to undergo cell death when treated with FlaTox or PrgJTox (Fig. 2C). In contrast, murine organoids underwent robust cell death in response to FlaTox or PrgJTox, as expected, indicating that these preparations of FlaTox and PrgJTox had the expected biological activity, but were not able to activate the inflammasome in primary or transformed human intestinal cells (Fig. 2C).

### Human intestinal epithelial cells express low levels of *NAIP* and *NLRC4* compared to human myeloid cells

Given the role of NAIP/NLRC4 in detecting and responding to *Salmonella* and bacterial T3SS ligands in murine IECs as well as murine and human macrophages (7– 12, 14, 16, 17), our findings that human IECs do not respond to T3SS ligands were surprising. Thus, we next asked if expression of *NAIP* and *NLRC4* in human IECs is comparable in Caco-2 cells, T84 cells, and human THP-1 macrophages (Fig. 3A, B).

**Figure 3:**
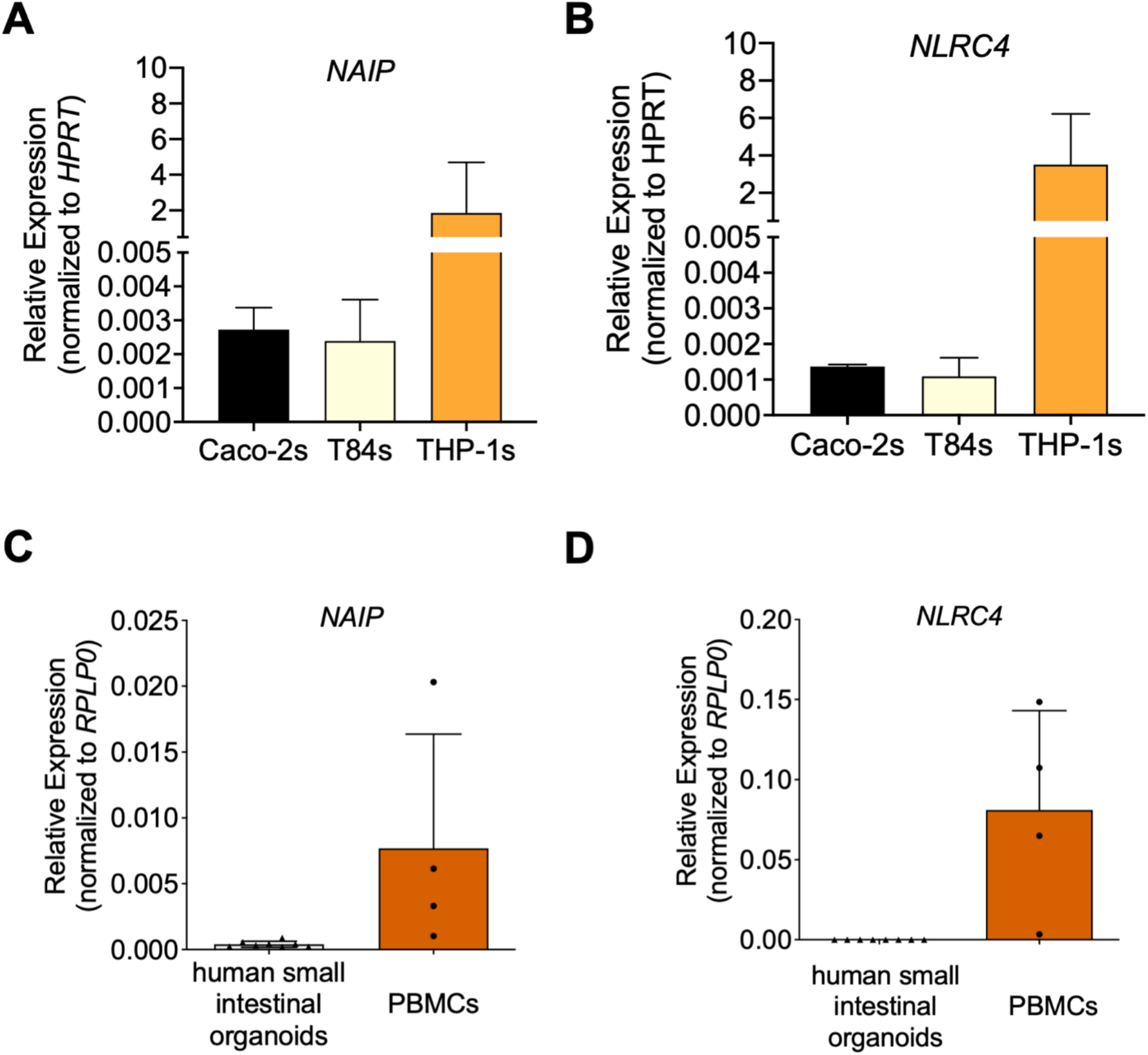
Human intestinal epithelial cells express low levels of NAIP/NLRC4 compared to human myeloid cells. Relative mRNA expression of *NAIP* and *NLRC4* compared to the housekeeping control *HPRT* or *RPLP0* as measured by qRT-PCR in (A, B) Caco-2 cells, T84 cells, and THP-1 macrophages, and in (C, D) human peripheral blood mononuclear cells (PBMCs) and human small intestinal organoids. Error bars represent the standard deviation of multiple wells. Data shown are representative of at least three independent experiments.

Consistent with their poor responsiveness to cytosolic delivery of NAIP ligands, both Caco-2 and T84 cells had very low expression of *NAIP* and *NLRC4* mRNA compared to human macrophages. Moreover, primary small intestinal organoids also expressed very low levels of *NAIP* and *NLRC4* compared to human PBMCs, indicating that this low expression was a general feature of human IECs (Fig. 3C, D). Collectively, these data suggest that human IECs express very low levels of NAIP and NLRC4, and this may partially explain why human IECs do not mount inflammasome responses to T3SS ligands.

### NAIP is not required for inflammasome activation in response to *Salmonella* infection of human intestinal cells

In murine macrophages and IECS as well as human macrophages, NAIP/NLRC4 contributes to inflammasome responses during *Salmonella* infection (11, 13, 14, 42). Even low levels of *NAIP* expression are sufficient to induce inflammasome responses in human macrophages (17). To formally test if the inflammasome activation we observe in human IECs infected with *Salmonella* (Fig. 1) requires NAIP, we used the Clustered Regularly Interspersed Palindromic Repeat (CRISPR) system, in conjunction with the RNA-guided exonuclease Cas9, to disrupt the *NAIP* gene in Caco-2 cells (Fig. S4). We sequenced two independent single cell clones each of *NAIP^-/-^* Caco-2 cells (*NAIP^-/-^* #7, *NAIP^-/-^* #8) to confirm appropriate targeting of *NAIP* in each line (Fig. S4). Caco-2 cells are polyploid, and we therefore found multiple mutant alleles for each CRISPR clone we sequenced (Fig. S4). For the two *NAIP^-/-^* clones, all of the changes resulted in a premature stop codon (Fig. S4).

To test whether inflammasome activation during *Salmonella* infection of human IECs requires NAIP, we infected WT or *NAIP^-/-^* Caco-2 cells with WT Stm or Δ*sipB* Stm and assayed for inflammasome activation by measuring IL-18 release and PI uptake (Fig. 4). As expected, WT Caco-2 cells infected with WT Stm released significantly increased levels of IL-18 and underwent cell death (Fig. 4A, 4B) and this response was dependent on the presence of the SPI-1 T3SS, as cells infected with Δ*sipB* Stm failed to undergo inflammasome activation (Fig. 4A, 4B). In *NAIP^-/-^* Caco-2 cells infected with WT Stm, we did not observe a decrease in inflammasome activation compared to WT Caco-2 cells (Fig. 4A, 4B). Both WT and *NAIP^-/-^* Caco-2 cells secreted similar levels of the inflammasome-independent cytokine IL-8 (Fig. 4C). Overall, these data indicate that NAIP is not required for inflammasome responses to SPI-1-expressing *Salmonella* in human IECs.

**Figure 4:**
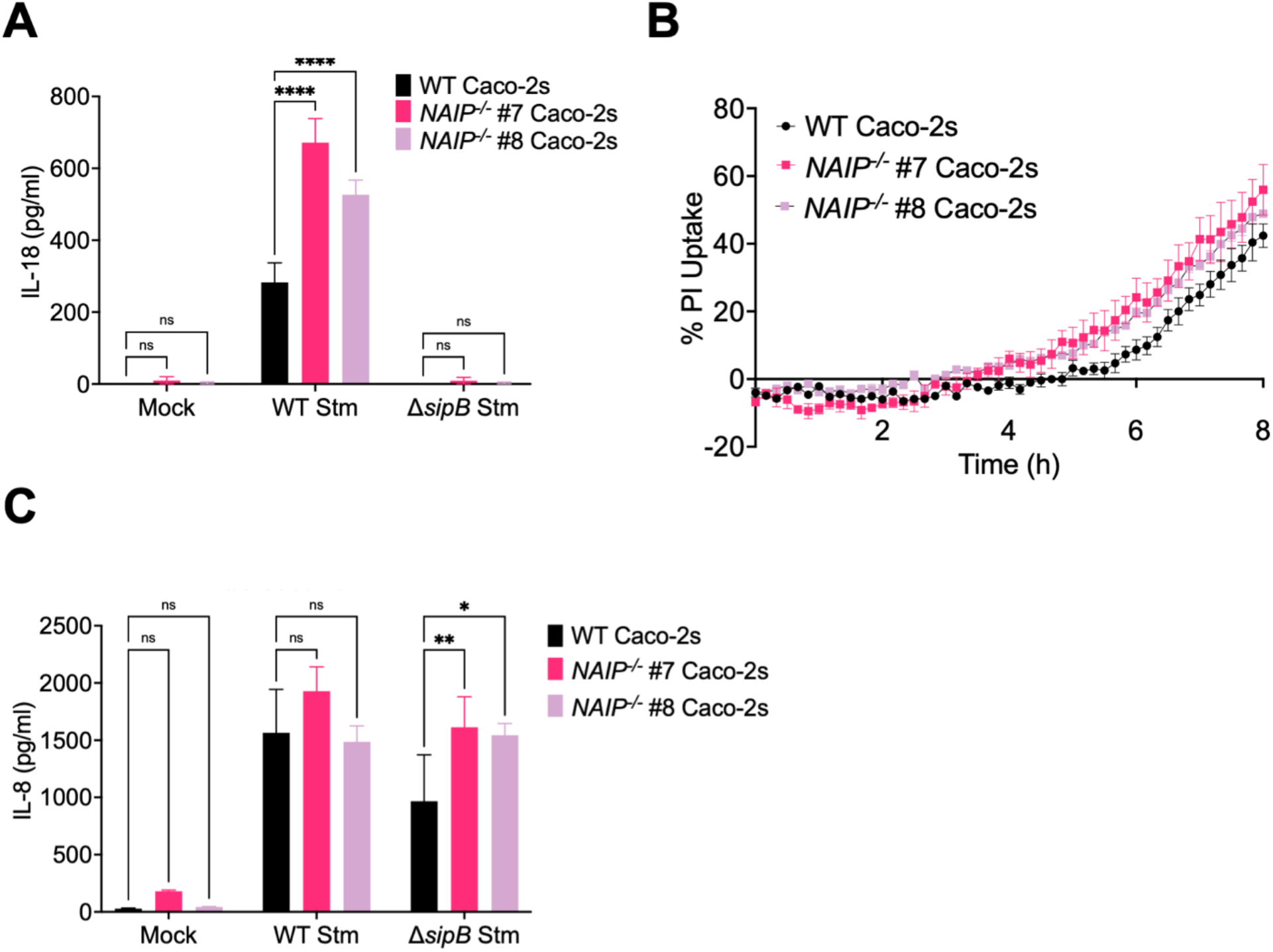
NAIP is not required for inflammasome responses to *Salmonella* in human intestinal epithelial cells. WT or two independent single cell clones of *NAIP^-/-^* Caco-2 cells were infected with PBS (Mock), WT *S*. Typhimurium, or Δ*sipB S*. Typhimurium for 6hrs. (A, C) Release of IL-18 or IL-8 into the supernatant was measured by ELISA. (B) Cell death was measured as percentage uptake of propidium iodide, normalized to cells treated with 1% Triton. (A, C) ns – not significant, * *p* < 0.05, ** *p* < 0.01, \*\*\*\*\**p* < 0.0001 by Dunnett’s multiple comparisons test. Error bars represent the standard deviation or the standard error of the mean (SEM) (B) of triplicate wells from one experiment. Data shown are representative of at least three independent experiments.

### NLRP3 and ASC are dispensable for inflammasome activation in response to *Salmonella* in human intestinal epithelial cells

Since the NAIP/NLRC4 inflammasome did not have a role in responding to *Salmonella* grown under SPI-1-inducing conditions in human IECs, we sought to determine whether another host cytosolic sensor could be responsible for inflammasome activation. One candidate is the NLRP3 inflammasome, which can be activated by a variety of stimuli during infection, including potassium efflux (18). In murine macrophages, the NLRP3 inflammasome is activated late during *Salmonella* infection (19). *Salmonella* infection also activates the NLRP3 inflammasome in human macrophages (26, 43). To determine if the inflammasome activation we observed in human IECs is dependent on the NLRP3 inflammasome, we infected WT Caco-2 cells that were pre-treated with MCC950, a potent chemical inhibitor of the NLRP3 inflammasome (44) and measured IL-18 release (Fig. 5A). As expected, cells treated with the DMSO control underwent robust inflammasome activation in response to WT Stm infection (Fig. 5A). Interestingly, cells treated with varying concentrations of MCC950 also exhibited similar levels of inflammasome activation as DMSO-control treated cells (Fig. 5A), suggesting that the NLRP3 inflammasome is not required for inflammasome responses to *Salmonella* in human IECs. We found that like *NAIP* and *NLRC4*, mRNA expression of *NLRP3* is also very low in Caco-2 cells compared to THP-1 macrophages (Fig. 5B), which may explain why NLRP3 inhibition does not prevent inflammasome activation in human IECs.

**Figure 5:**
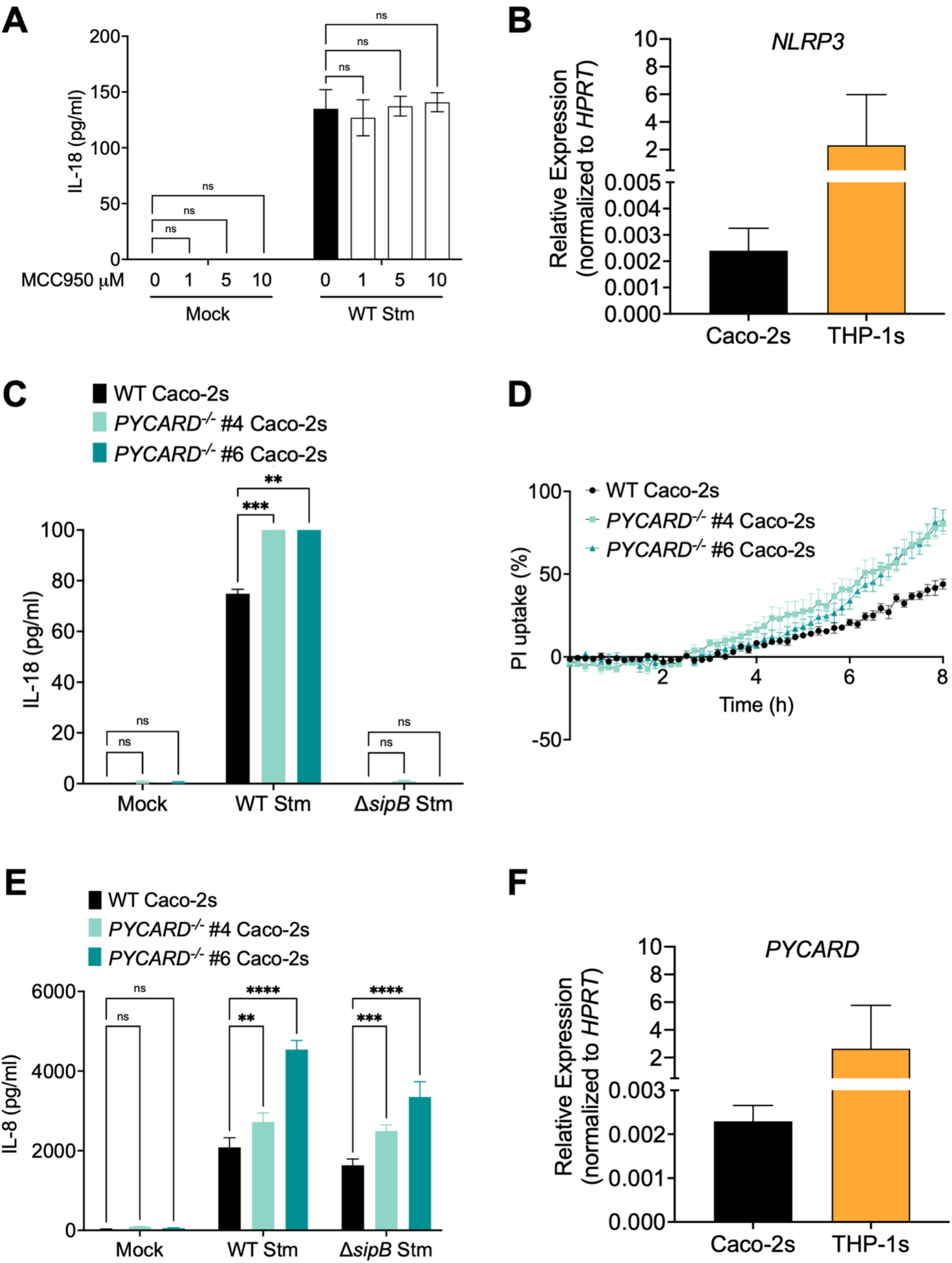
NLRP3 and ASC are dispensable for inflammasome responses to *Salmonella* in human intestinal epithelial cells. (A) WT Caco-2 cells were primed for 3hrs with 400 ng/ml of Pam3CSK4. One hour prior to infection, cells were treated with the indicated concentrations of MCC950 or DMSO as a vehicle control. Cells were then infected with PBS (Mock) or WT *S*. Typhimurium for 6hrs. Release of IL-18 into the supernatant was measured by ELISA. (B, F) Relative mRNA expression of *NLRP3* and *PYCARD* compared to the housekeeping control *HPRT* as measured by qRT-PCR in Caco-2 cells and THP-1 macrophages. (C, D, E) WT or two independent single cell clones of *PYCARD^-/-^* Caco-2 cells were infected with PBS (Mock), WT *S*. Typhimurium, or Δ*sipB S*. Typhimurium. (C, D) Release of IL-18 or IL-8 into the supernatant were measured by ELISA at 6hpi. (E) Cell death was measured as percentage uptake of propidium iodide, normalized to cells treated with 1% Triton. (A, C, E) ns – not significant, ** *p* < 0.01, *** *p* < 0.001, **** *p* < 0.0001 by Dunnett’s multiple comparisons test. Error bars represent the standard deviation or the standard error of the mean (SEM) (D) of triplicate wells from one experiment. Data shown are representative of at least three independent experiments.

In addition to the NAIP/NLRC4 and NLRP3 inflammasomes, there are many other inflammasomes that can get activated in response to bacterial infections. The majority of these inflammasomes, including AIM2, IFI16, NLRP3, NLRP6, NLRP7, and pyrin, use an adaptor protein called ASC to recruit and activate downstream caspases (4). To determine if ASC-dependent inflammasomes participate in the response to *Salmonella* in human IECs, we tested if ASC is required for inflammasome activation. Using CRISPR/Cas9, we disrupted *PYCARD*, the gene that encodes for ASC, in Caco-2 cells (Fig. S5). We sequence-validated two independent single cell clones of *PYCARD^-/-^* Caco-2 cells (*PYCARD^-/-^* #4, *PYCARD^-/-^* #6) (Fig. S5). Most of the mutations resulted in a premature stop codon or the mutated protein bore no resemblance to the WT protein sequence (Fig. S5). mRNA expression of *PYCARD* was also abrogated in the KO clones relative to WT Caco-2 cells (Fig. S6A). We did not detect any ASC protein expression even in WT Caco-2 cells (Fig. S6B).

We next infected WT or *PYCARD^-/-^* Caco-2 cells with WT Stm or Δ*sipB* Stm and assayed for inflammasome activation (Fig. 5C, D). As expected, WT Caco-2 cells infected with WT Stm released significantly increased levels of IL-18 and underwent cell death (Fig. 5C, D). This was dependent on the presence of the SPI-1 T3SS, as cells infected with Δ*sipB* Stm failed to undergo inflammasome activation (Fig. 5C, D). In *PYCARD*^-/-^ Caco-2 cells infected with WT Stm, we did not observe a decrease in inflammasome activation compared to WT Caco-2 cells, indicating that ASC is dispensable for IL-18 release and cell death in Caco-2 cells in response to Stm. Both WT and *PYCARD^-/-^* Caco-2 cells exhibited similar levels of IL-8 release, an inflammasome-independent cytokine (Fig. 5E). In addition, expression of *PYCARD* mRNA in Caco-2 cells was very low compared to THP-1 macrophages (Fig. 5F). Collectively, these data indicate that NLRP3 and ASC-dependent inflammasomes are not required for inflammasome responses to *Salmonella* infection in human IECs.

### Caspase-1 is partially required for inflammasome activation in response to *Salmonella* in human intestinal epithelial cells

Inflammasomes recruit various caspases which can then cleave and process IL-1 and IL-18 cytokines and mediate pyroptosis (45). For example, the murine NAIP/NLRC4 inflammasome can recruit both caspase-1 and caspase-8 in response to *Salmonella* infection of murine IECs (14). In human macrophages, NAIP/NLRC4, NLRP3, caspase-1, and caspase-8 are recruited to the same macromolecular complex during *Salmonella* infection (26). Notably, although expression of *CASP1* mRNA in Caco-2 cells is lower than that in THP-1 macrophages (Fig. 6A), *CASP1* expression in Caco-2 cells is still higher than expression of other inflammasome genes we have assessed so far (compare axes in Fig. 6A to Fig. 3A, 3B, 5B, 5F). Similarly, expression of *CASP1* in small intestinal organoids, although lower than that observed in human PBMCs, is higher than expression of other inflammasome-related genes (compare axes in Fig. 6B to Fig. 3C, 3D). This suggests that caspase-1 may be an important contributor to inflammasome responses in IECs.

**Figure 6:**
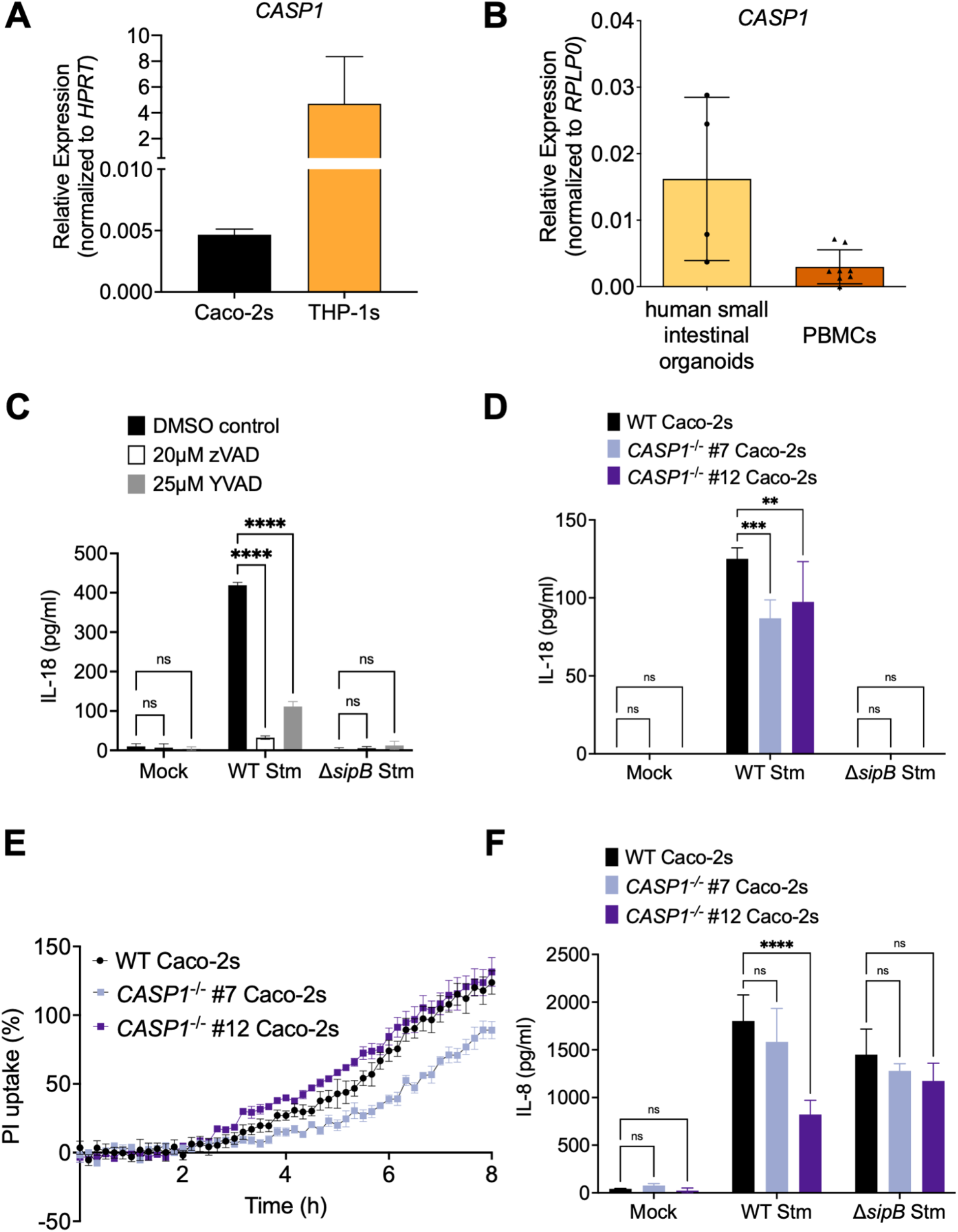
Caspase-1 is partially required for inflammasome responses to *Salmonella* in human intestinal epithelial cells. (A, B) Relative mRNA expression of *CASP1* compared to the housekeeping control *HPRT* as measured by qRT-PCR in Caco-2 cells, THP-1 macrophages, human peripheral blood mononuclear cells (PBMCs), and human small intestinal organoids. (C) WT Caco-2 cells were primed with 400 ng/ml of Pam3CSK4 for 3h. One hour prior to infection, cells were treated with 20 μM of pan-caspase inhibitor Z-VAD(OMe)-FMK, 25 μM of caspase-1 inhibitor Ac-YVAD-cmk, or DMSO as a vehicle control. Cells were then infected with PBS (Mock), WT *S*. Typhimurium, or Δ*sipB S*. Typhimurium for 6hrs. Release of IL-18 into the supernatant was measured by ELISA. (D – F) WT or two independent single cell clones of *CASP1^-/-^* Caco-2 cells were infected with PBS (Mock), WT *S*. Typhimurium, or Δ*sipB S*. Typhimurium. (D, F) Release of IL-18 or IL-8 into the supernatant were measured by ELISA at 6hpi. (E) Cell death was measured as percentage uptake of propidium iodide, normalized to cells treated with 1% Triton. (C, D, F) ns – not significant, ** *p* < 0.01, **** *p* < 0.0001 by Dunnett’s multiple comparisons test. Error bars represent the standard deviation or the standard error of the mean (SEM) (E) of triplicate wells from one experiment. Data shown are representative of at least three independent experiments.

To test the contribution of caspases to inflammasome responses during *Salmonella* infection in human IECs, we pretreated Caco-2 cells with pharmacological inhibitors targeting caspases of interest (ZVAD: pan-caspase inhibitor or YVAD: inhibitor for caspase-1). We then infected cells with WT Stm or Δ*sipB* Stm and assayed for inflammasome activation by measuring IL-18 release (Fig. 6C). DMSO-treated cells infected with WT Stm released IL-18, whereas cells treated with the inhibitors had a significant defect in IL-18 release. Treatment with ZVAD, the pan-caspase inhibitor, resulted in a lower level of IL-18 release compared to treatment with the caspase-1 inhibitor YVAD, suggesting that in addition to caspase-1, other caspases are also important during *Salmonella* infection of human IECs. As expected, cells infected with Δ*sipB* Stm demonstrated no inflammasome activation regardless of inhibitor treatment. These results suggest that caspase-1 contributes to inflammasome responses to *Salmonella* in human IECs.

Since pharmacological inhibitors preferentially targeting individual caspases can have cross-reactivity with other caspases, we used CRISPR/Cas9 to disrupt *CASP1* in Caco-2 cells (Fig. S7, S8). We sequenced and validated two independent single cell clones of *CASP1^-/-^* Caco-2 cells (*CASP1^-/-^* #7, *CASP1^-/-^* #12) (Fig. S7, S8). All mutations resulted in a premature stop codon (Fig. S7, S8). We further confirmed decreased mRNA expression of *CASP1* in *CASP1^-/-^* #7 Caco-2 cells (Fig. S7B). We were unable to detect protein expression in both WT Caco-2 cells and *CASP1^-/-^* #7 Caco-2 cells by western blot (Fig. S7C), indicating that caspase-1 is poorly expressed in Caco-2 cells.

To test whether inflammasome activation during *Salmonella* infection requires caspase-1 in human IECs, we infected WT or *CASP1^-/-^* Caco-2 cells with WT or Δ*sipB* Stm and assayed for subsequent inflammasome activation (Fig. 6D, E). As expected, WT Caco-2 cells infected with WT Stm released significant levels of IL-18 and underwent cell death (Fig. 6D, E). This response was dependent on the presence of the SPI-1 T3SS, as cells infected with Δ*sipB* Stm failed to undergo inflammasome activation (Fig. 6D). Consistent with YVAD inhibitor treatment, *CASP1*^-/-^ Caco-2 cells infected with WT Stm showed a statistically significant decrease in, but not complete abrogation of, IL-18 release at 6hpi (Fig. 6D). There was also a slight delay in uptake of PI in *CASP1^-/-^* #7 Caco-2 cells relative to WT Caco-2 cells (Fig. 6E). In contrast, both WT and KO Caco-2 cells exhibited similar levels of IL-8 release, an inflammasome-independent cytokine (Fig. 6F). Overall, these data suggest that caspase-1 is partially required for the inflammasome response to *Salmonella* infection in human IECs.

### Caspase-4 is required for inflammasome responses to *Salmonella* in human intestinal epithelial cells

In mice, in addition to caspase-1, a second caspase, caspase-11, responds to *Salmonella* infection (19, 21). Caspase-11 and its human orthologs caspases-4/5 detect cytosolic LPS and form the noncanonical inflammasome (24, 25). Caspases-4/5 are important for late inflammasome responses to *Salmonella* infection of human intestinal epithelial cells, specifically when *Salmonella* escapes to the cytoplasm from its *Salmonella*-containing vacuole (SCV) (20). To test the contribution of the caspase-4/5 inflammasome earlier during infection with *Salmonella* grown under SPI-1-inducing conditions, we transfected Caco-2 cells with siRNAs targeting *CASP4, CASP5* or both. We then infected the cells with WT or Δ*sipB* Stm and assayed for inflammasome activation by measuring IL-18 release (Fig. 7A). As expected, cells treated with a control scrambled siRNA exhibited IL-18 secretion upon infection with WT Stm, but failed to undergo IL-18 secretion when infected with Δ*sipB* Stm (Fig. 7A). However, knockdown of *CASP4*, either alone, or in conjunction with *CASP5*, resulted in nearly complete abrogation of IL-18 secretion in cells infected with WT Stm compared to control siRNA-treated cells, suggesting that caspase-4 is required for inflammasome activation (Fig. 7A). Knockdown of *CASP5* alone resulted in a partial and significant decrease in IL-18 secretion, indicating that while caspase-5 may be playing a role, it is not absolutely required (Fig. 7A). We observed moderately high (67 – 75%) siRNA-mediated knockdown efficiencies (Fig. S9A). Release of the inflammasome-independent cytokine IL-8 was comparable across conditions (Fig. S9B).

**Figure 7:**
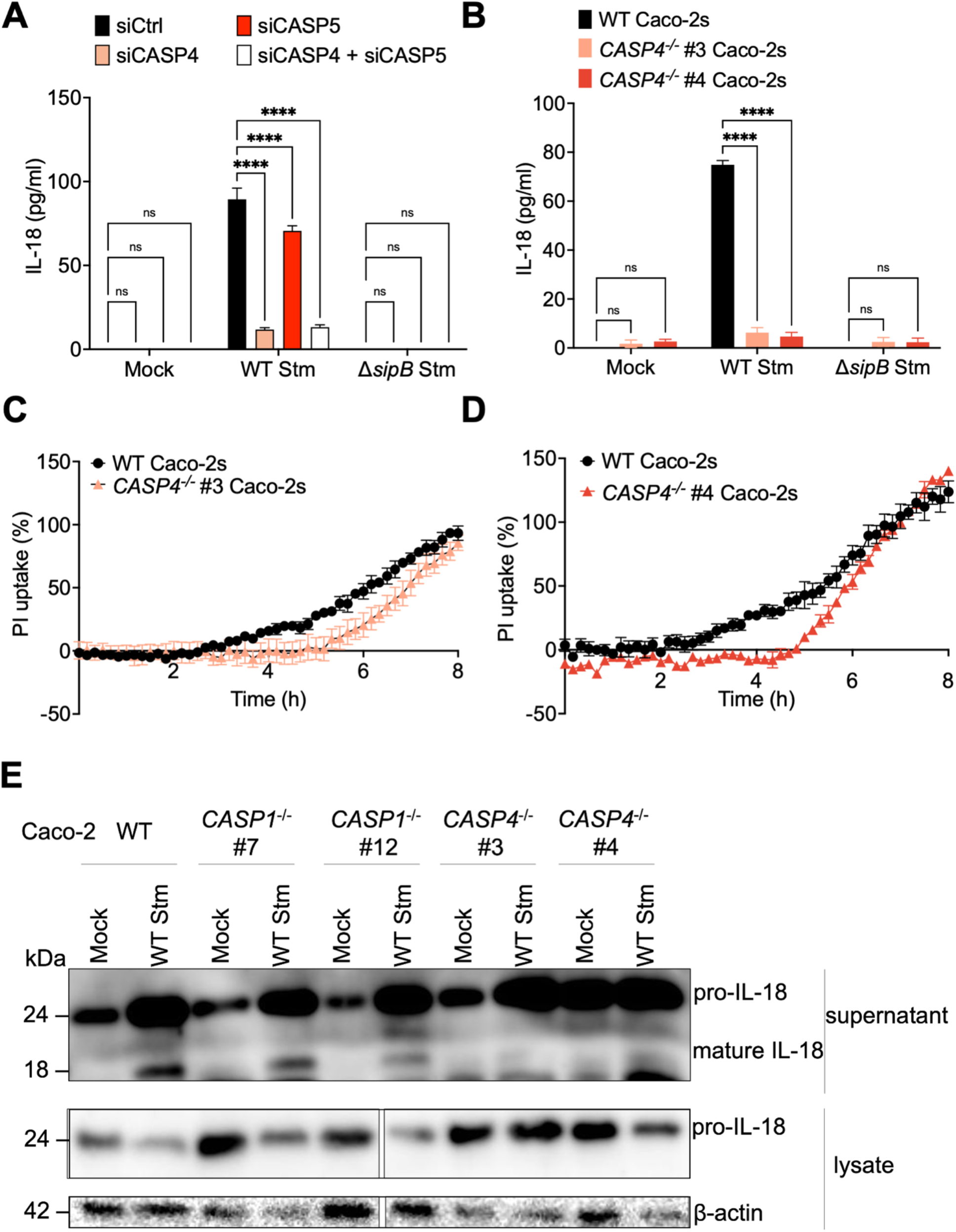
Caspase-4 is required for inflammasome responses to *Salmonella* in human intestinal epithelial cells. (A) WT Caco-2 cells were treated with siRNA targeting *CASP4, CASP5,* or a control scrambled siRNA for 72h. Cells were primed with 400 ng/ml of Pam3CSK4 for 3hrs. Cells were then infected with PBS (Mock), WT *S*. Typhimurium, or Δ*sipB S*. Typhimurium for 6hrs. Release of IL-18 was measured by ELISA (B – E) WT or two independent clones of *CASP4^-/-^* or *CASP1^-/-^* (E only) Caco-2 cells were infected with PBS (Mock), WT *S*. Typhimurium, or Δ*sipB S*. Typhimurium for 6hrs. (B) Release of IL-18 into the supernatant was measured by ELISA at 6hpi. (C, D) Cell death was measured as percentage uptake of propidium iodide, normalized to cells treated with 1% Triton. (E) Lysates and supernatants collected 6hpi were immunoblotted for IL-18 and β-actin. (A, B) ns – not significant, **** *p* < 0.0001 by Tukey’s (A) or Dunnett’s (B) multiple comparisons test. Error bars represent the standard deviation or the standard error of the mean (SEM) (C, D) of triplicate wells from one experiment. Data shown are representative of at least three independent experiments.

To definitively test the requirement of caspase-4 in early inflammasome responses to SPI-1-expressing *Salmonella*, we disrupted *CASP4* in Caco-2 cells using CRISPR/Cas9 (Fig. S10, S11). We sequenced and validated two independent single cell clones of *CASP4^-/-^* Caco-2 cells (*CASP4^-/-^* #3, *CASP4^-/-^* #4) (Fig. S10). All mutations resulted in a premature stop codon (Fig. S10). Both clones exhibited decreased mRNA expression of *CASP4* and no protein expression relative to WT Caco-2 cells (Fig. S11).

We then infected WT or *CASP4^-/-^* Caco-2 cells with WT or Δ*sipB* Stm and assayed for subsequent IL-18 secretion and cell death as readouts for inflammasome activation (Fig. 7B – E). As expected, WT Caco-2 cells infected with WT Stm released significant levels of cleaved IL-18 and underwent cell death (Fig. 7B – E), whereas cells infected with Δ*sipB* Stm failed to release IL-18 (Fig. 7B). Consistent with our findings with siRNA knockdown of *CASP4*, *CASP4^-/-^* Caco-2 cells infected with WT Stm failed to release IL-18 and exhibited a delay in PI uptake at 4-6 hpi (Fig. 7B – E). Interestingly, *CASP4^-/-^* Caco-2 cells began to undergo cell death later in infection, suggesting that while caspase-4 is required for inducing cell death early in infection, it may not be absolutely required at later timepoints of infection. Like WT cells, *CASP4^-/-^* cells still released substantial levels of the inflammasome-independent cytokine IL-8 following infection (Fig. S9C). Collectively, these data suggest that caspase-4 is required for early inflammasome responses to SPI-1-expressing *Salmonella* infection in human IECs.

## Discussion

In this study, we have demonstrated that Caco-2 cells undergo inflammasome activation in response to early infection with SPI-1-expressing *Salmonella* that requires the presence of the SPI-1 T3SS, and that GSDMD-mediated pore formation is required for release of IL-18 and cell death (Fig. 1, S1). Unexpectedly however, individual bacterial T3SS ligands or flagellin were not sufficient to induce inflammasome activation in Caco-2 cells or intestinal organoids (Fig. 2, S2, S3). Additionally, we found that neither the NAIP nor NLRP3 canonical inflammasomes (Fig. 4, S4, 5), are required for inflammasome responses during *Salmonella* infection, which may be due to low mRNA expression of *NAIP*, *NLRC4*, and *NLRP3* in human IECs and human intestinal organoids (Fig. 3, 5). Moreover, we found that ASC, a shared adaptor protein that several inflammasomes use to recruit caspases, is also not required for inflammasome activation (Fig. 5, S5, S6), suggesting that inflammasomes that require ASC for their function are likely not playing a role in human IECs during *Salmonella* infection.

Interestingly, we found that caspase-1 is partially required for inflammasome activation in human IECs (Fig. 6, S7, S8), leaving the unanswered question of what upstream sensor is activating caspase-1. Finally, we found that caspase-4 is necessary for early inflammasome responses to SPI-1-expressing *Salmonella* (Fig. 7, S9 – S11).

NAIP/NLRC4 inflammasome activation in IECs is critical for control of *Salmonella* infection in mice (13–15). It was therefore surprising that NAIP was dispensable in human IECs during *Salmonella* infection (Fig. 3, S3 – S6). Perhaps a lack of robust NAIP/NLRC4 inflammasome responses in human IECs partially underlies why some intestinal bacterial pathogens with T3SSs, such as *Shigella flexneri,* enterohemorrhagic *Escherichia coli,* and enteropathogenic *E. coli,* can cause disease in humans but not mice. Indeed, mice are normally resistant to *S. flexneri,* but mice lacking the NAIP/NLRC4 inflammasome can be robustly colonized by *S. flexneri* and develop a disease resembling human shigellosis (46).

The lack of NAIP activation may be due to low expression of *NAIP* and *NLRC4* mRNA in human IEC lines as well as human small intestinal organoids (Fig. 3). Single cell transcriptome analysis of cells intestinal cells from human donors demonstrate that only a small subset of intestinal cells express *NAIP* at detectable levels, while *NLRC4* expression is below the limit of detection (47). However, it is possible that a population of IECs exist that express *NAIP* and *NLRC4* at low levels that may be below the limit of detection of single-cell transcriptome analysis, but may be sufficient to activate the inflammasome in response to infection under physiological conditions. Indeed, single cell transcriptome analysis of murine small intestines reveal that expression of *NLRC4* is also very low in murine IECs (48), but murine IECs undergo robust NAIP/NLRC4 inflammasome activation despite this relatively low expression (13–15). Despite our findings that NAIP is dispensable in Caco-2 cells to respond to *Salmonella* infection and human intestinal organoids do not mount inflammasome responses to T3SS ligands, it is possible that under physiological conditions, this inflammasome still plays an important role during bacterial infection or disease pathogenesis. Indeed, human patients with activating NLRC4 mutations exhibit gastrointestinal symptoms (49–52).

A recent study comparing inflammasome responses of human and murine IECs during *Salmonella* infection reported that while caspase-1 was required for inflammasome activation in murine IECs, it was dispensable in late inflammasome responses to *Salmonella* in human IECs (22). We found caspase-1 to be partially required for inflammasome activation in IECs infected with *Salmonella* (Fig. 6, S7, S8). Differences in our experimental conditions likely explain this difference in findings. Our *Salmonella* was grown under SPI-1-inducing conditions, and we assayed for inflammasome activation at a slightly earlier timepoint. Perhaps caspase-1 plays a role in early inflammasome responses to SPI-1-expressing *Salmonella* in human IECs, and it is less important at later timepoints when *Salmonella* has shifted to expressing its SPI-2 T3SS.

The identity of the host sensor that is acting upstream of caspase-1 in human IECs remains unknown. Humans have 22 NLRs that could serve as potential cytosolic sensors of bacterial structures and activity (53). Most of these NLRs do not contain a CARD domain, and they would therefore require an adaptor protein such as ASC to recruit caspases. Given our finding that ASC is not required for inflammasome activation in human IECs (Fig. 5, S5, S6), it is likely that the putative host sensor upstream of caspase-1 contains its own CARD domain. Alternatively, it may interact with a different CARD domain-containing adaptor protein than ASC. Future studies will focus on identifying host factors that different caspases interact with during *Salmonella* infection of human IECs.

In agreement with previous studies (20, 22), we found caspase-4 to be required for inflammasome activation in response to *Salmonella* in human IECs (Fig 7, S9 – S11). Importantly, caspase-4 activation restricts bacterial replication, and specifically restricts replication of a subpopulation of cytosolic *Salmonella* (20, 22). It is unclear whether *Salmonella*’s access into the cytosol is mediated by the bacteria or host, and future studies will explore the mechanisms by which cytosolic populations of *Salmonella* arise and how they influence inflammasome activation in human IECs.

In mice, inflammasome activation mediates control of *Salmonella* by restricting bacterial replication, extruding infected cells, and preventing systemic dissemination of *Salmonella* (13, 15, 20). Human IECs have been shown to undergo extrusion, but whether this occurs in a caspase-4-dependent manner remains unknown (54). While we know that human IECs and enteroids restrict bacterial replication in a caspase-4-dependent manner (20, 22), it remains unknown if inflammasome activation results in other mechanisms of control of infection. In mice, IL-18 release during *Salmonella* infection recruits natural killer (NK) cells that are critical for early mucosal inflammatory responses (55). We observe robust release of the inflammasome-dependent cytokine IL-18, but the downstream role of this cytokine in the human intestine during *Salmonella* infection has not been explored. Additionally, while our study focused exclusively on *Salmonella* infection, it is worth exploring if enteric pathogens with similar lifestyles to *Salmonella*, such as *Shigella*, or ones with different lifestyles, such as the extracellular pathogen *Yersinia*, elicit similar responses. Future studies that interrogate downstream consequences of inflammasome activation in human IECs in response to various enteric pathogens could shed light on human mucosal inflammatory responses to bacterial pathogens.

Overall, our data indicate that *Salmonella* infection of human IECs triggers inflammasome pathways that are distinct from that in mice. Pathways that are activated in mice and important for control of infection, such as the NAIP and NLRP3 inflammasomes, were unexpectedly not required for inflammasome responses in human IECs under the conditions we investigated. Instead, inflammasome responses in human IECs required caspases-1 and −4. Our findings provide a foundation for future studies aimed at uncovering the relative contribution of different caspases, and the downstream responses that they mediate in human IECs.

## Acknowledgements

We thank members of Igor Brodsky’s and Sunny Shin’s laboratories for scientific discussion. We thank Meghan Wynosky-Dolfi for technical advice. We thank Russell Vance, Randilea Nichols, and Jeannette Tenthorey for providing anthrax toxin-based reagents, and JD Sauer for providing the *Listeria* strains and constructs for generating ActA fusion proteins. We thank Jared Fisher for providing spheroids of human duodenum and colon, and William Messer and Zoe Lyski for providing cDNA of human PBMCs. Work in the Shin laboratory is supported by NIH/NIAID grants AI118861 and AI123243, and the Linda Pechenik Montague Investigator Award from the University of Pennsylvania Perelman School of Medicine. Work in the Brodsky laboratory is supported by NIH/NIAID grants AI128520 and AI139102. S.S. and I.B. are both recipients of the Burroughs-Wellcome Fund Investigators in the Pathogenesis of Infectious Disease Award. I.R. is supported by OHSU start-up funding. N.N. is a recipient of the American Heart Association Predoctoral Fellowship 19PRE34380315. R.B. was supported by the FWF-Immunity in Cancer & Allergy PhD program of the University of Salzburg and the Austrian Marshall Plan Foundation Scholarship. J.Z. is a recipient of the NIH/NIAID Microbial Pathogenesis and Genomics training grant 5T32AI141393-03.

