## Supplemental Information for "*Salmonella* Typhimurium induces NAIP/NLRC4- and NLRP3/ASC-independent, caspase-1/4-dependent inflammasome activation in human intestinal epithelial cells"

### Supplemental Materials and Methods

#### Cell Culture of C2Bbe1 cells

C2Bbe1 cells (CRL-2102; ATCC) were maintained in DMEM supplemented with 0.01mg/ml human transferrin, 10% (vol/vol) heat-inactivated FBS, 100 IU/mL penicillin, and 100 μg/mL streptomycin. Cells were maintained at 37°C in a humidified incubator. Cells were replated and assays conducted as described for other intestinal cell lines in the main body.

#### Polarization of intestinal epithelial cells

Transparent membrane (PET) 1 μm pore size cell culture inserts (Greiner Bio-One 662610) in a 24-well plate were coated with collagen coating solution containing 30 μg/ml collagen, 10 μg/ml fibronectin, and 10 μg/ml BSA in DMEM and incubated overnight at 37°C. Caco-2 cells were plated in growth medium containing Corning MITO+ serum extender (Fisher Scientific CB-50006). After 24 hours, the growth medium was replaced with Corning enterocyte differentiation medium (Fisher Scientific 355357) containing MITO+ serum extender. The media was replaced daily and after three days of incubation in differentiation medium, the transepithelial electrical resistance was measured using an Epithelial Volt/Ohm (TEER) Meter (World Precision Instruments) to ensure that it is ≥250 $\Omega.cm^2$ prior to infection or treatments. All treatments were administered on the apical side of the cells.

Anthrax toxin-mediated delivery of bacterial ligands into human intestinal epithelial cell lines

Cells were primed with 400 ng/ml Pam3CSK4 (Invivogen) for 4 hours prior to treatment. Recombinant proteins (PA and LFn-PrgJ) were kindly provided by Russell Vance (1). PA and LFn doses for *in vitro* delivery were: 4 μg/ml PA and 8 ng/ml LFn-PrgJ. Cells were incubated at 37°C for 16 hours.

Generation of CRISPR Cas9 knockouts in Caco-2 cells

To knockout genes in Caco-2 cells, plasmids encoding the desired guide RNA (gRNA) and Cas9 in the pLentiCRISPR v2 plasmid were purchased from GenScript. The sequences of these gRNAs are listed below:

| | | |
|---|---|---|
| *NAIP* | gRNA 1 | : ACATTGCCAAGTACGACATA |
| *PYCARD* | gRNA 1 | : CATGTCGCGCAGCACGTTAG |
| *CASP1* | gRNA 1 | : GACAGTATTCCTAGAAGAAC |
| *CASP4* | gRNA 1 | : TCCTGCAGCTCATCCGAATA |

Paul Bates at the University of Pennsylvania kindly provided pCMV-VSV-G and psPAX2 plasmids to produce lentiviral particles. Lentiviral particles were generated in HEK293T cells. HEK293T cells were plated at $2 \times 10^6$ cells per 10cm dish in 10mL of DMEM supplemented with 10% (vol/vol) heat-inactivated FBS, 2 mM L-glutamine, 100 IU/mL penicillin, and 100 μg/mL streptomycin. After 24 hours, plasmids were transfected using Lipofectamine 2000 according to manufacturer's instructions. 1 μg of pCMV-VSV-G, 2.5 μg of psPAX2, and 5 μg of pLentiCRISPR v2 encoding the appropriate gRNA, and 50μL of Lipofectamine 2000 was used per dish. Transfected HEK293T cells were incubated for 18 hours at 37°C, and the media was then aspirated and replaced with 6mL of fresh growth media. After an additional 24 hours, the supernatant containing lentiviral particles was harvested and filtered using a 0.22µM filter. 5 x $10^5$ Caco-2 cells were infected in 1mL of virus-containing media with 8 µg/mL of polybrene in a 12-well plate. The cells were then spun at 1250 × *g* for 90 minutes at 30°C, and subsequently incubated at 37°C for 24 hours. After 24 hours, the virus-containing media was replaced with Caco-2 growth media and incubated at 37°C. After an additional 24 hours, puromycin was added to a final concentration of 10 µg/mL. The cells were maintained in puromycin for approximately 4 weeks and gradually expanded from a 12-well plate to larger dishes, ending in a T175 flask. Cells were then harvested for clonal selection.

For clonal selection, cells were plated in 96-well plates at 0.5 cell per well or 2 cells per well in 200µl of growth media and were incubated for 4-8 weeks until single clones were visible in the bottom of the well. 10-12 single clones for each gene were selected and gradually expanded from a 96-well plate to larger dishes, ending in a T175 flask. Cells were then plated in 24-well plates at a concentration of 3 × $10^5$ cells per well in 500µl of media and harvested for purification of DNA for validation.

Validation of CRISPR Cas9 Caco-2 single clones for knockouts

DNA was harvested from cells and purified using the DNeasy Blood and Tissue kit (Qiagen). The following primers were used to amplify the genomic region containing the gRNA target sequence via PCR:

*NAIP* forward : 5'- CCGTACAGCTCATGGATACCACAG -3'

*NAIP* reverse : 5'- GTACCTGTAAAGACAAAGCCAGCC -3'

*PYCARD* forward : 5’- GACCTCACCGACAAGCTG -3’

*PYCARD* reverse : 5’- GGGGTAGGAGGAACAGAAAG -3’

*CASP1* forward : 5’- GGGCATTTGCAATGTCCATGCA -3’

*CASP1* reverse : 5’- CCAGGCTTGTGCTGCATGAC -3’

*CASP4* forward : 5’- GGAAAGGCCAAATTTAACCCCAAC -3’

*CASP4* reverse : 5’- ATGGTTACTGTCATCCCCACCC -3’

The PCR product was purified using the PCR Cleanup kit (Qiagen). A poly A-tail was added to the purified PCR product by adding together 7µl of PCR product, 5 Units of Taq DNA polymerase, 1X PCR Buffer containing $MgCl_2$, and 0.2mM dATP. The reaction was then incubated at 70°C for 30 minutes. 2µL of this product was then ligated into the pGEM-T Vector System and transformed into DH5α competent cells according to manufacturer’s instructions (A1360; Promega). Blue-white screening was used to select approximately 15 colonies that were PCR-screened using the same primers shown above and run on an agarose gel to confirm that they contained the insert. At least 10 positive colonies were sequenced using one of the following primers:

M13/pUC forward : 5’- CCCAGTCACGACGTTGTAAAACG -3’

M13/pUC reverse : 5’- AGCGGATAACAATTTCACACAGG -3’

<u>Knockdown efficiency of siRNA-mediated knockdown of genes</u>

To calculate knock-down efficiency, RNA was extracted and cDNA synthesized as described in main body. The following primers from PrimerBank (identification numbers within parenthesis) (2–4) were used:

*CASP4* (73622124c1) Forward: 5’- CAAGAGAAGCAACGTATGGCA -3’

| | | |
|---|---|---|
| | Reverse: | 5'- AGGCAGATGGTCAAACTCTGTA -3' |
| *CASP5* (209870072c2) | Forward: | 5'- TTCAACACCACATAACGTGTCC -3' |
| | Reverse: | 5'- GTCAAGGTTGCTCGTTCTATGG -3' |

mRNA levels siRNA-treated cells were normalized to housekeeping gene *HPRT* and control siRNA-treated cells using the $2^{-\Delta\Delta CT}$ (cycle threshold) method (5).

<u>CRISPR/Cas9 immunoblot analysis</u>

Cell lysates were harvested for immunoblot analysis by adding 1X SDS/PAGE sample buffer to cells and boiled for 5 minutes. Samples were separated by SDS/PAGE on a 12% (vol/vol) acrylamide gel, and transferred to PVDF Immobilon-P membranes (Millipore). Primary antibodies specific for human caspase-1 (2225S; Cell Signaling), human caspase-4 (4450S; Cell Signaling), human ASC (D086-3, MBL International) and β-actin (4967L; Cell Signaling), and HRP-conjugated secondary antibody anti-rabbit IgG (7074S; Cell Signaling) were used. ECL Western Blotting Substrate or SuperSignal West Femto (both from Pierce Thermo Scientific) were used as the HRP substrate for detection.

Supplemental Figures

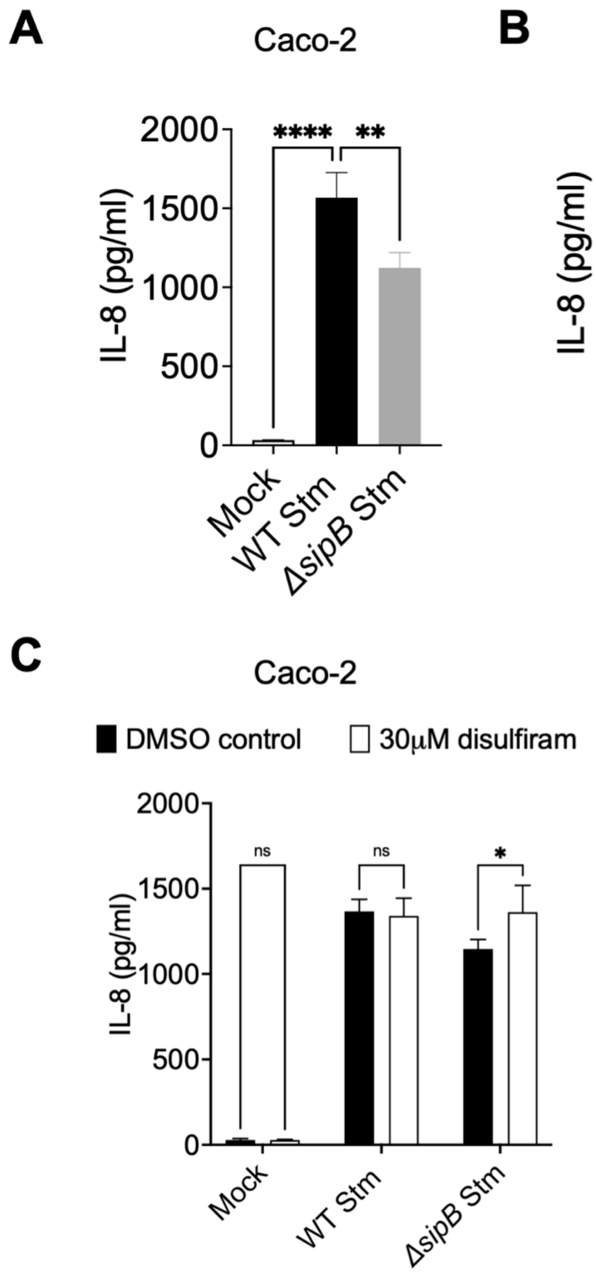


**Figure S1 (related to Figure 1): *Salmonella* infection induces IL-8 release in human intestinal epithelial cells.** (A) Caco-2 cells or (B) T84 cells were infected with PBS (Mock), WT *S*. Typhimurium, or Δ*sipB S*. Typhimurium. Release of IL-8 into the supernatant was measured by ELISA at 6hpi. (C) Caco-2 cells were treated with 30 μM disulfiram or DMSO as a vehicle control 1 hour prior to infection. Cells were then infected with PBS (Mock), WT *S.* Typhimurium, or Δ*sipB S*. Typhimurium. Release of IL-8 into the supernatant was measured by ELISA at 6hpi. ns – not significant, * $p < 0.05$, ** $p < 0.01$, **** $p < 0.0001$ by Dunnett's multiple comparisons test (A), or by unpaired t-test (B), or by Šídák's multiple comparisons (C). Error bars represent the standard deviation of triplicate wells from one experiment. Data shown are representative of at least three independent experiments.

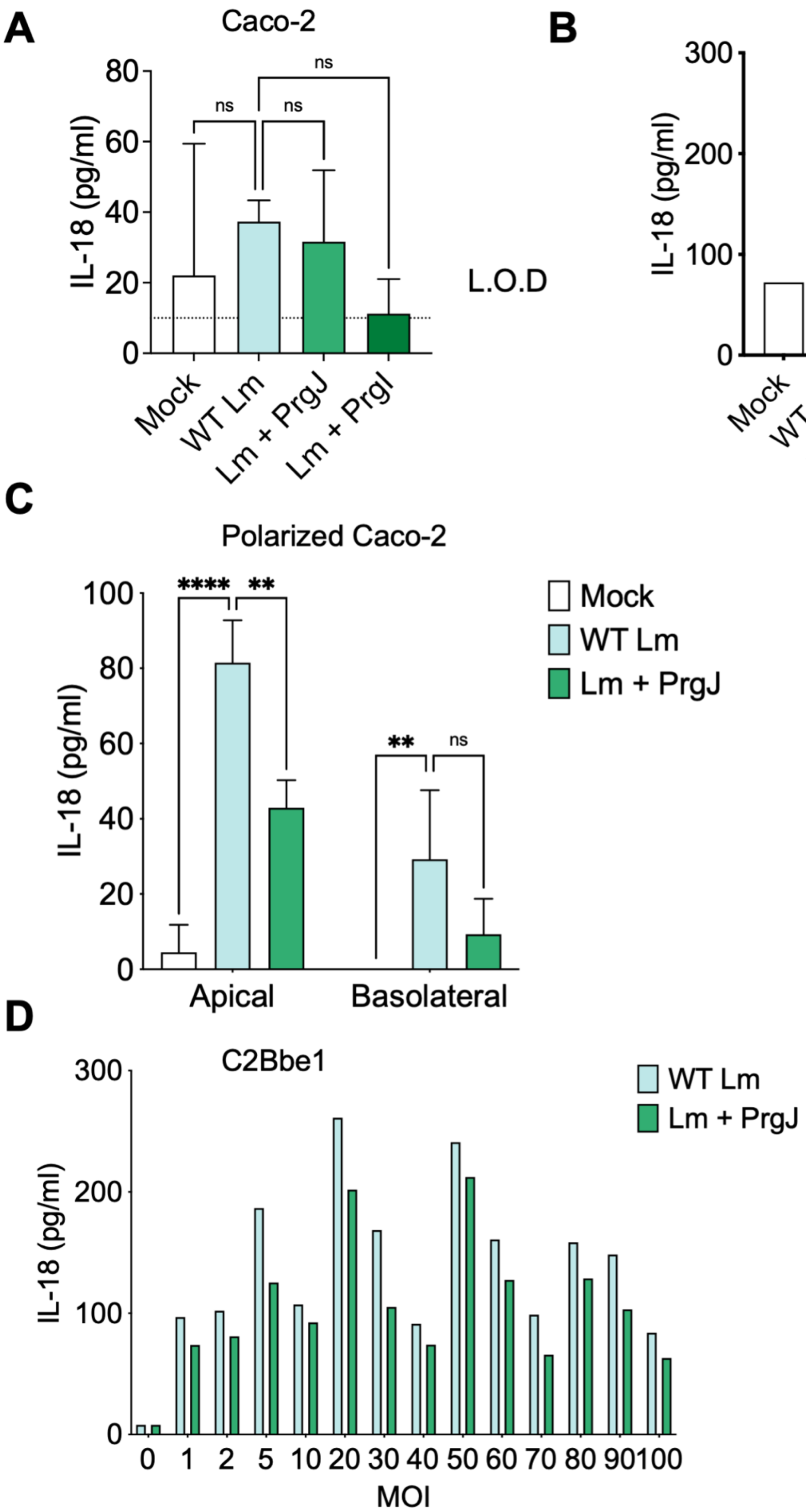
A
Caco-2
IL-18 (pg/ml)
ns
ns
ns
L.O.D
Mock
WT Lm
Lm + PrgJ
Lm + PrgI
B
T84
IL-18 (pg/ml)
Mock
WT Lm
Lm + PrgI
C
Polarized Caco-2
IL-18 (pg/ml)
****
**
**
ns
Mock
WT Lm
Lm + PrgJ
Apical
Basolateral
D
C2Bbe1
IL-18 (pg/ml)
WT Lm
Lm + PrgJ
MOI

**Figure S2 (related to Figure 2): Bacterial T3SS ligands delivered using *Listeria* do not activate the inflammasome in human intestinal epithelial cells.** WT Caco-2 cells (A), T84 cells (B), polarized Caco-2 cells (C) or C2Bbe1 cells (D) were primed for 3h with 400 ng/ml of Pam3CSK4 and infected with PBS (Mock), WT *L. monocytogenes* (WT Lm), or *L. monocytogenes* expressing *S.* Typhimurium SPI-1 inner rod (Lm + PrgJ), or SPI-1 needle (Lm + PrgI) at the following MOIs: Caco-2 = 5; T84 = 25; polarized Caco-2 = 100; C2Bbe1 = as indicated in graph. Release of IL-18 into the supernatant was measured by ELISA at 16hpi. ns – not significant, ** $p < 0.01$, **** $p < 0.0001$ by Dunnett's multiple comparisons test. L.O.D indicates the limit of detection of the assay. Error bars represent the standard deviation of triplicate wells from one experiment. (B) and (D) were conducted in duplicate wells.

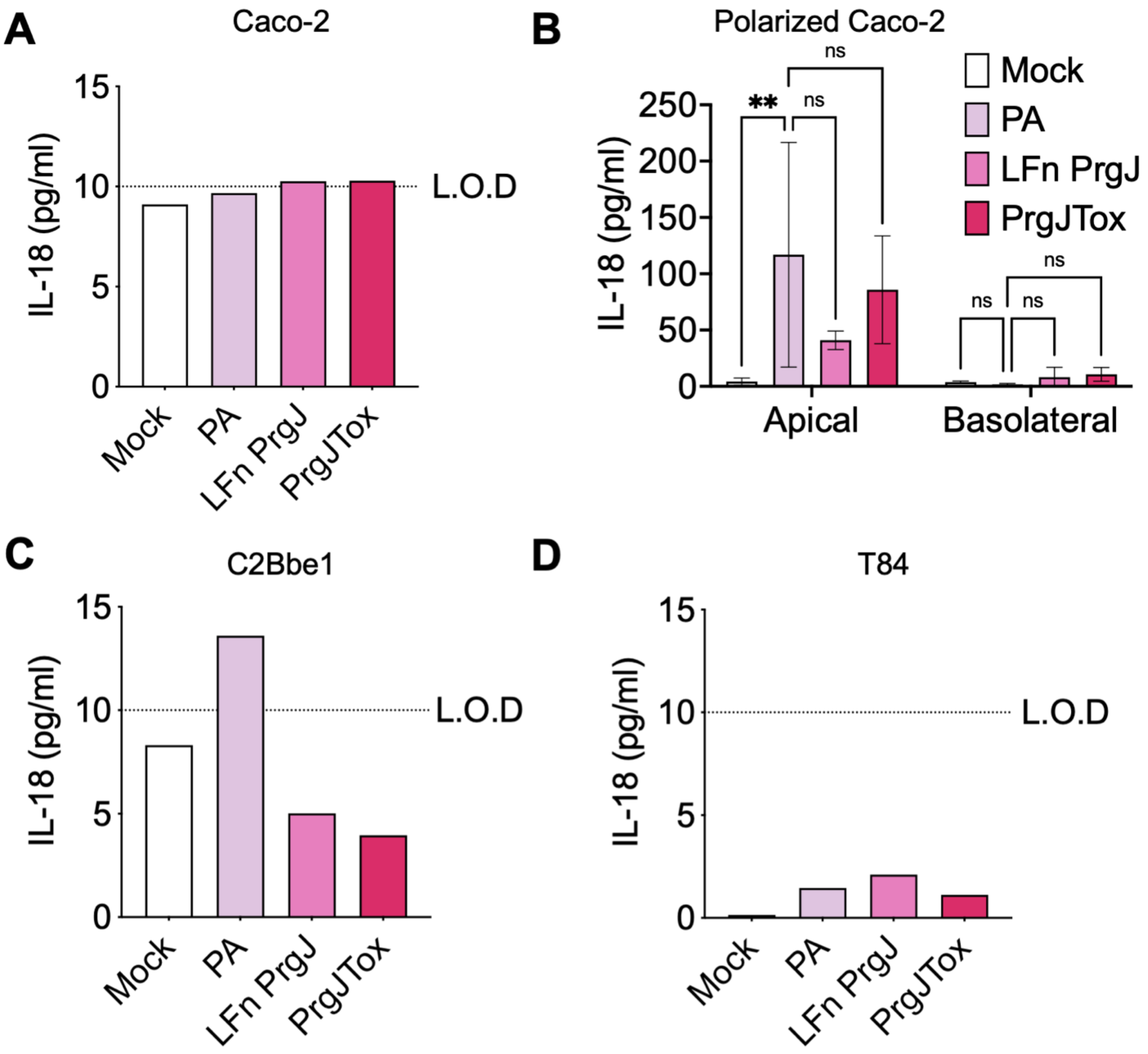


**Figure S3 (related to Figure 2): Bacterial T3SS ligands delivered using the anthrax toxin system do not activate the inflammasome in human intestinal epithelial cells.** WT Caco-2 cells (A), polarized Caco-2 cells (B), C2Bbe1 cells (C), or T84 cells (D) were primed for 3h with 400 ng/ml of Pam3CSK4. Cells were then treated with PBS (Mock), PA alone, LFnPrgJ alone, or PA+LFnPrgJ (PrgJTox) for 16 hours. Release of IL-18 into the supernatant was measured by ELISA. For polarized Caco-2s (B), supernatants were collected from both apical and basolateral compartments. L.O.D indicates the limit of detection of the assay. Error bars represent the standard deviation of triplicate wells from one experiment. (A, C, D) were conducted in duplicate wells. Data shown are from one independent experiment (A, D) or are representative of two independent experiments (B, C). ns – not significant, ** $p < 0.01$ by Dunnett's multiple comparisons test.

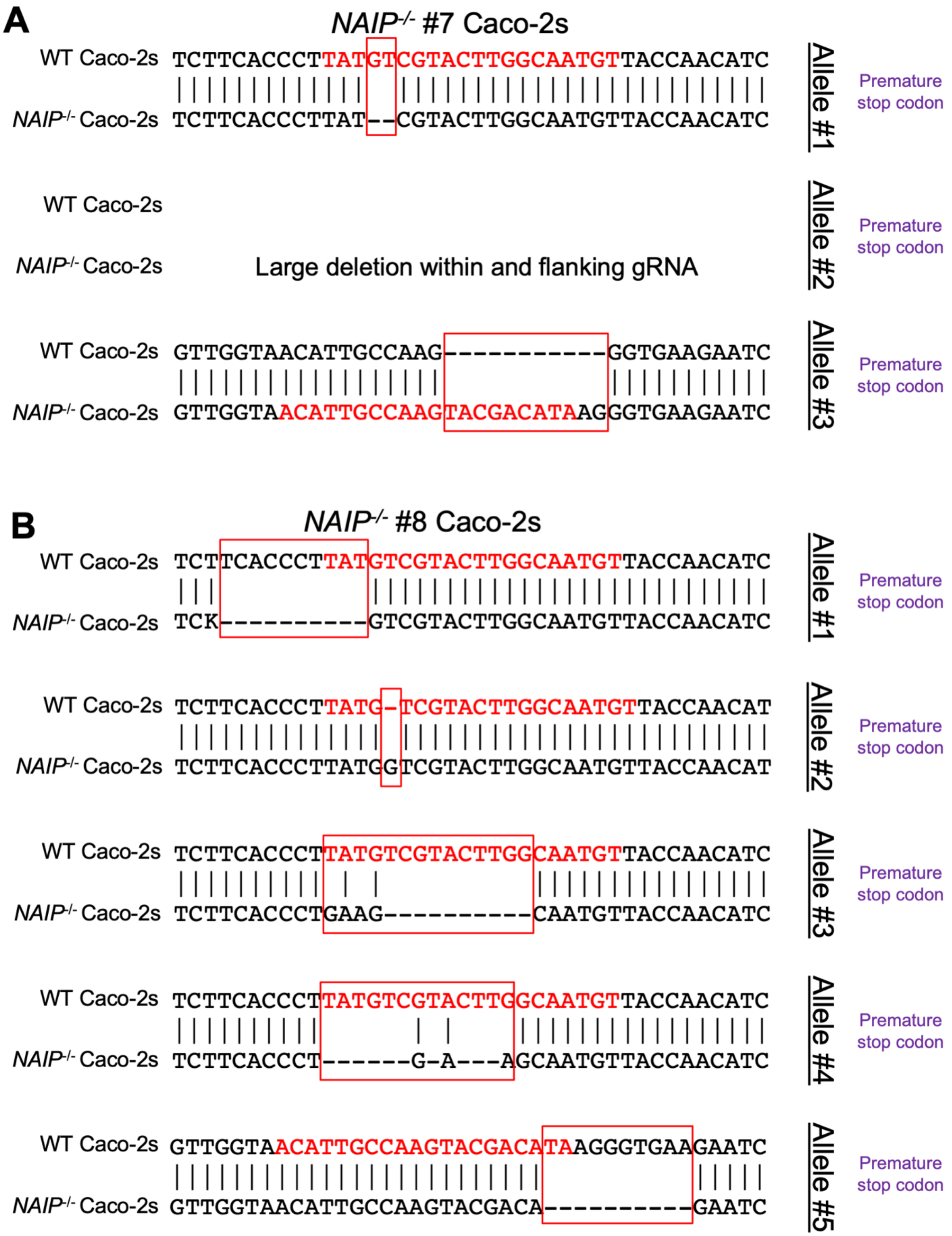
A
NAIP-/- #7 Caco-2s
WT Caco-2s TCTTCACCCTTATGTCGTACTTGGCAATGTTACCAACATC
NAIP-/- Caco-2s TCTTCACCCTTAT--CGTACTTGGCAATGTTACCAACATC
Allele #1
Premature stop codon
WT Caco-2s
NAIP-/- Caco-2s
Large deletion within and flanking gRNA
Allele #2
Premature stop codon
WT Caco-2s GTTGGTAACATTGCCAAG-----------GGTGAAGAATC
NAIP-/- Caco-2s GTTGGTAACATTGCCAAGTACGACATAAGGGTGAAGAATC
Allele #3
Premature stop codon
B
NAIP-/- #8 Caco-2s
WT Caco-2s TCTTCACCCTTATGTCGTACTTGGCAATGTTACCAACATC
NAIP-/- Caco-2s TCK----------GTCGTACTTGGCAATGTTACCAACATC
Allele #1
Premature stop codon
WT Caco-2s TCTTCACCCTTATG-TCGTACTTGGCAATGTTACCAACAT
NAIP-/- Caco-2s TCTTCACCCTTATGGTCGTACTTGGCAATGTTACCAACAT
Allele #2
Premature stop codon
WT Caco-2s TCTTCACCCTTATGTCGTACTTGGCAATGTTACCAACATC
NAIP-/- Caco-2s TCTTCACCCTGAAG----------CAATGTTACCAACATC
Allele #3
Premature stop codon
WT Caco-2s TCTTCACCCTTATGTCGTACTTGGCAATGTTACCAACATC
NAIP-/- Caco-2s TCTTCACCCT------G-A---AGCAATGTTACCAACATC
Allele #4
Premature stop codon
WT Caco-2s GTTGGTAACATTGCCAAGTACGACATAAGGGTGAAGAATC
NAIP-/- Caco-2s GTTGGTAACATTGCCAAGTACGACA----------GAATC
Allele #5
Premature stop codon

**Figure S4 (related to Figure 4): Validation of *NAIP* mutant Caco-2 cells generated with CRISPR/Cas9 genome editing.** Sequence alignments of WT Caco-2, *NAIP*$^{-/-}$ #7 Caco-2, and *NAIP*$^{-/-}$ #8 Caco-2 are shown for multiple alleles. Red text represents the target guide RNA sequence and red boxes represent the mutated region. Purple text represents the predicted impact of the mutation on the amino acid sequence.

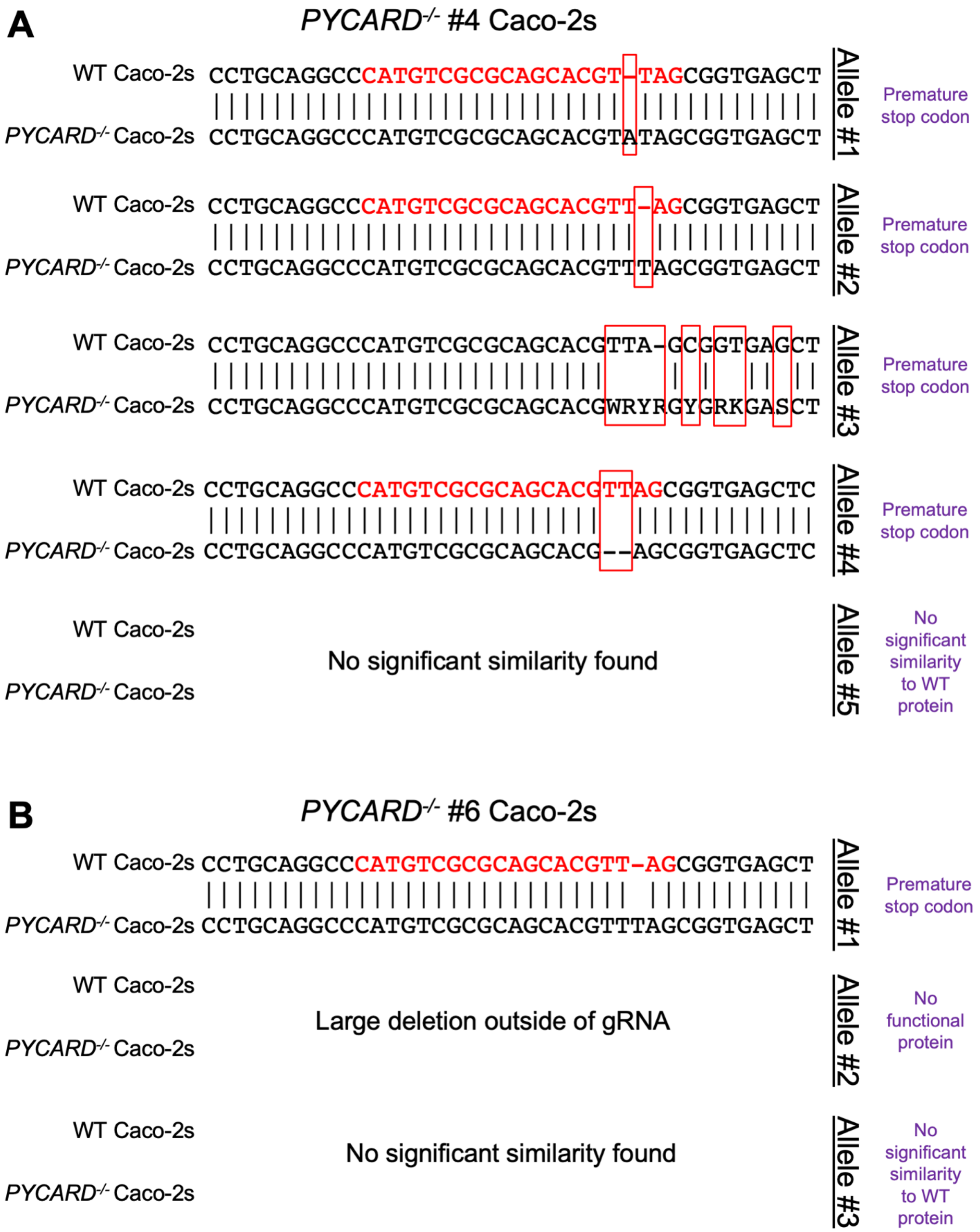


**Figure S5 (related to Figure 5): Validation of *PYCARD* mutant Caco-2 cells generated with CRISPR/Cas9 genome editing.** Sequence alignments of WT Caco-2,

*PYCARD*$^{-/-}$ #4, and *PYCARD*$^{-/-}$ #6 Caco-2 are shown for multiple alleles. Red text represents the target guide RNA sequence and red boxes represent the mutated region. Purple text represents the predicted impact of the mutation on the amino acid sequence.

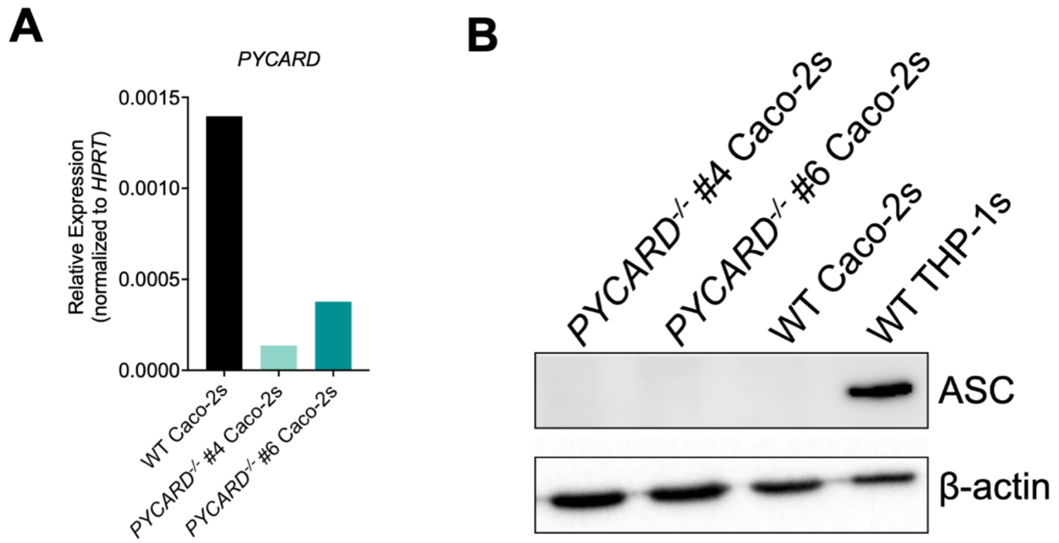


**Figure S6 (related to Figure 5): Expression of ASC in *PYCARD* mutant Caco-2 cells generated with CRISPR/Cas9 genome editing.** (A) Relative mRNA expression of *PYCARD* compared to the housekeeping control *HPRT* as measured by qRT-PCR in WT and *PYCARD*$^{-/-}$ #4 Caco-2s and *PYCARD*$^{-/-}$ #6 Caco-2s. (B) WT THP-1s, WT Caco-2s, *PYCARD*$^{-/-}$ #4 Caco-2s, and *PYCARD*$^{-/-}$ #6 Caco-2s lysates were immunoblotted for ASC and β-actin.

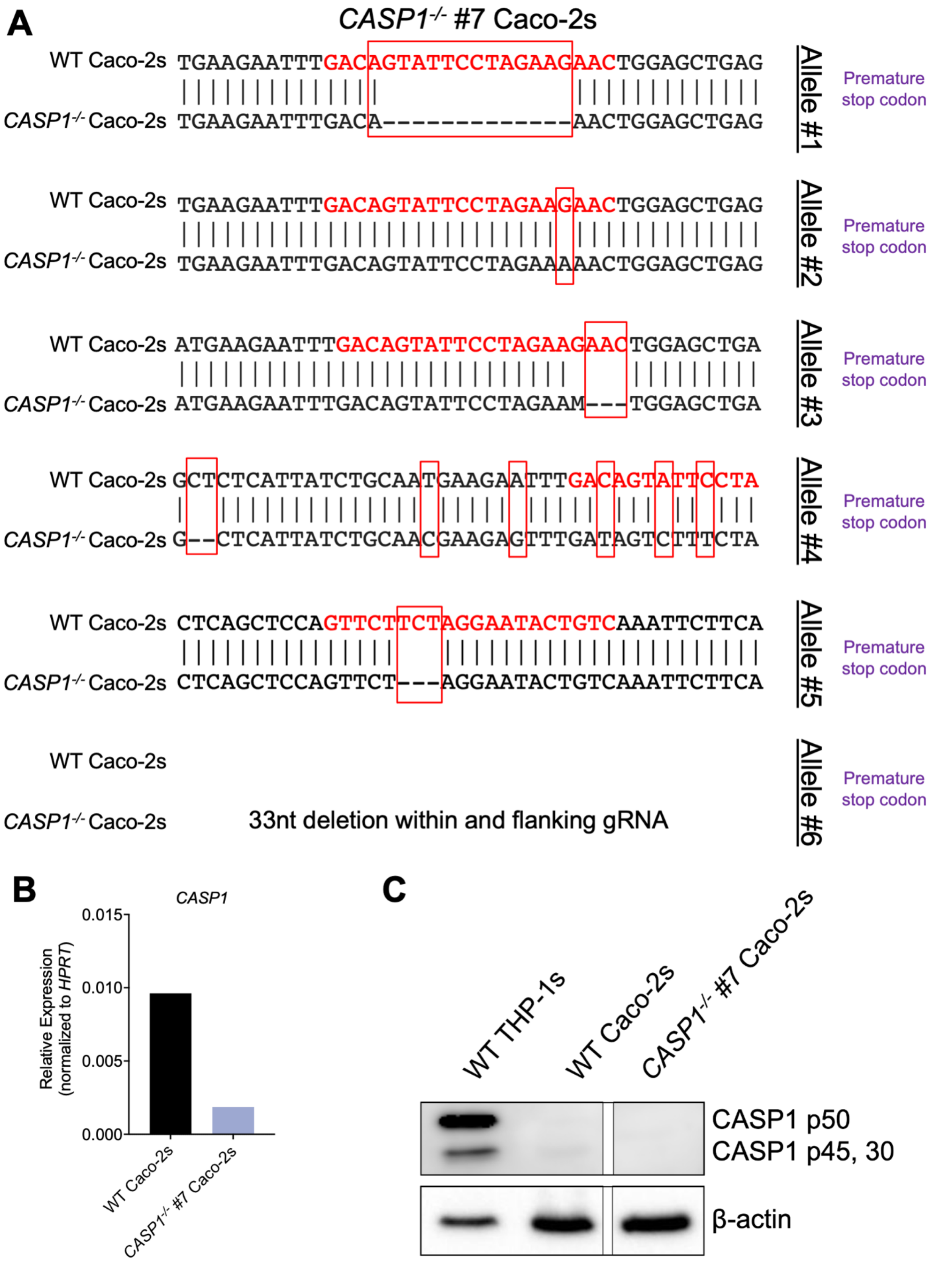
A
CASP1-/- #7 Caco-2s
WT Caco-2s TGAAGAATTTGACAGTATTCCTAGAAGAACTGGAGCTGAG
CASP1-/- Caco-2s TGAAGAATTTGACA-------------AACTGGAGCTGAG
Allele #1
Premature stop codon
WT Caco-2s TGAAGAATTTGACAGTATTCCTAGAAGAACTGGAGCTGAG
CASP1-/- Caco-2s TGAAGAATTTGACAGTATTCCTAGAAAAACTGGAGCTGAG
Allele #2
Premature stop codon
WT Caco-2s ATGAAGAATTTGACAGTATTCCTAGAAGAACTGGAGCTGA
CASP1-/- Caco-2s ATGAAGAATTTGACAGTATTCCTAGAAM---TGGAGCTGA
Allele #3
Premature stop codon
WT Caco-2s GCTCTCATTATCTGCAATGAAGAATTTGACAGTATTCCTA
CASP1-/- Caco-2s G--CTCATTATCTGCAACGAAGAGTTTGATAGTCTTTCTA
Allele #4
Premature stop codon
WT Caco-2s CTCAGCTCCAGTTCTTCTAGGAATACTGTCAAATTCTTCA
CASP1-/- Caco-2s CTCAGCTCCAGTTCT---AGGAATACTGTCAAATTCTTCA
Allele #5
Premature stop codon
WT Caco-2s
CASP1-/- Caco-2s
33nt deletion within and flanking gRNA
Allele #6
Premature stop codon
B
CASP1
Relative Expression (normalized to HPRT)
0.015
0.010
0.005
0.000
WT Caco-2s
CASP1-/- #7 Caco-2s
C
WT THP-1s
WT Caco-2s
CASP1-/- #7 Caco-2s
CASP1 p50
CASP1 p45, 30
β-actin

**Figure S7 (related to Figure 6): Validation of *CASP1* mutant Caco-2 Clone 7 generated with CRISPR/Cas9 genome editing.** (A) Sequence alignments of WT Caco-2s, and *CASP1*$^{-/-}$ #7 Caco-2s are shown for multiple alleles. Red text represents the target guide RNA sequence and red boxes represent the mutated region. Purple text represents the predicted impact of the mutation on the amino acid sequence. (B) Relative mRNA expression of *CASP1* compared to the housekeeping control *HPRT* in WT Caco-2s and *CASP1*$^{-/-}$ #7 Caco-2s, as measured by qRT-PCR. (C) WT THP-1s, WT Caco-2s, *CASP1-/-* #7 Caco-2 lysates were immunoblotted for ASC and β-actin.

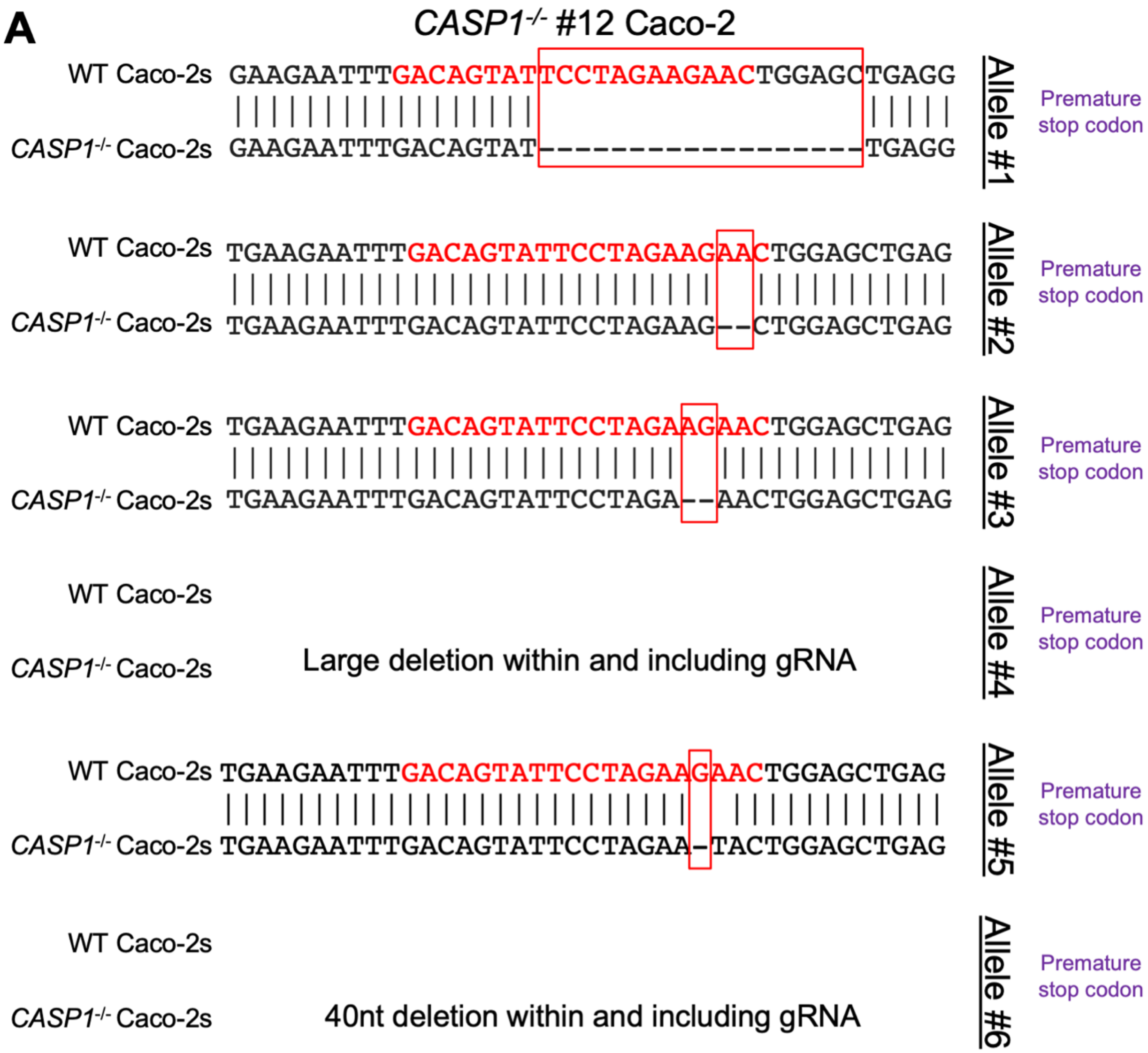

**Figure S8 (related to Figure 6): Validation of *CASP1* mutant Caco-2 Clone 12 generated with CRISPR/Cas9 genome editing.** Sequence alignments of WT Caco-2, and *CASP1*$^{-/-}$ #12 Caco-2 are shown for multiple alleles. Red text represents the target guide RNA sequence and red boxes represent the mutated region. Purple text represents the predicted impact of the mutation on the amino acid sequence.

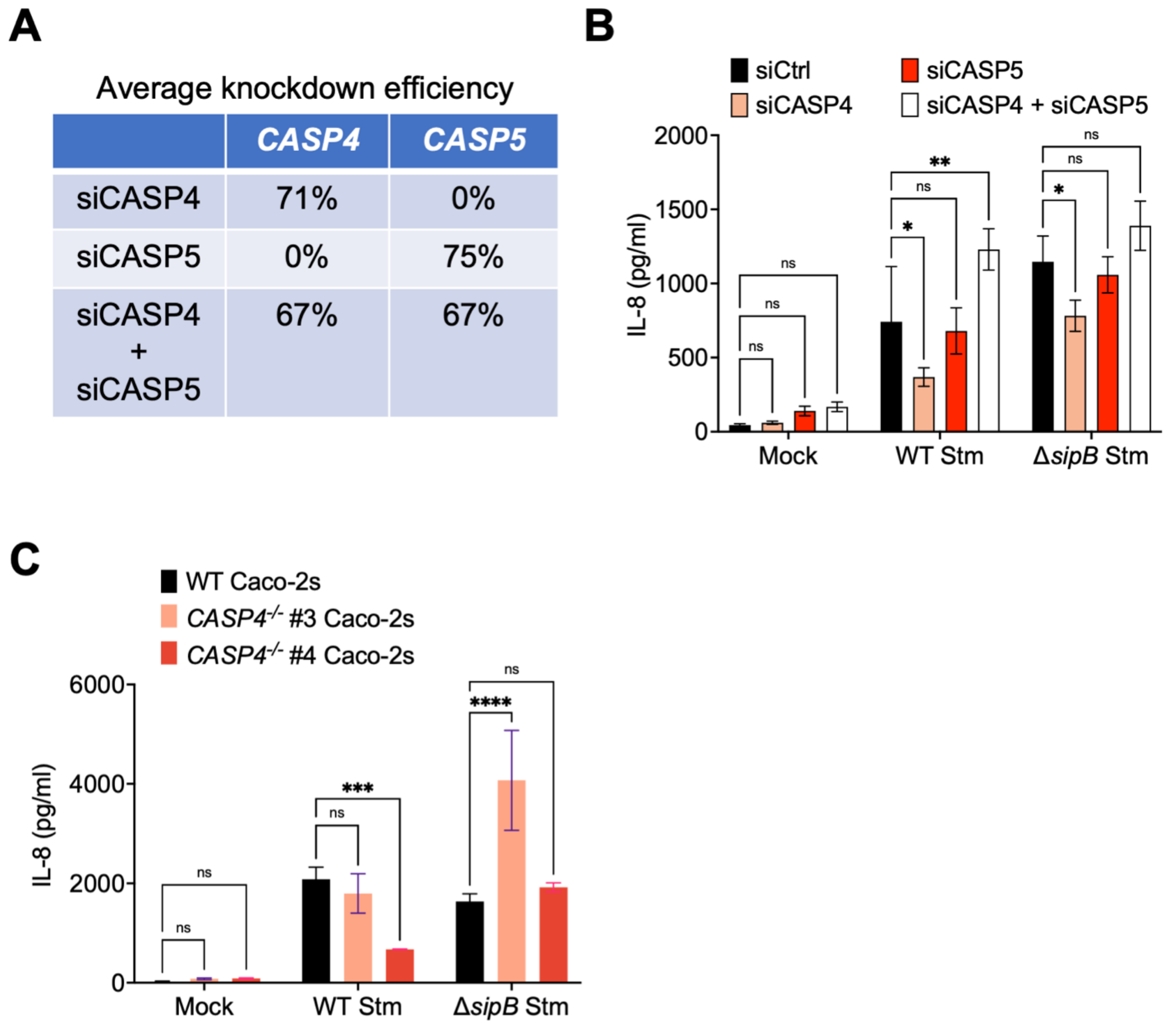


**Figure S9 (related to Figure 7): *Salmonella* infection induces CASP4/5-dependent inflammasome activation in human intestinal epithelial cells.** (A, B) WT Caco-2 cells were treated with siRNA targeting *CASP4, CASP5,* or a control scrambled siRNA for 72h. Cells were primed with 400 ng/ml of Pam3CSK4 for 3hrs. Cells were then infected with PBS (Mock), WT *S*. Typhimurium*,* or Δ*sipB S*. Typhimurium for 6hrs. (A) Knockdown efficiency was measured by qRT-PCR and normalized to housekeeping gene *HPRT*, and calculated relative to control-siRNA-treated cells. (B) Release of IL-8 into the supernatant was measured by ELISA at 6hpi. (C) WT or two independent clones of *CASP4*$^{-/-}$ Caco-2 cells were infected with PBS (Mock), WT *S*. Typhimurium, or Δ*sipB S*. Typhimurium for 6hrs. Release of IL-8 into the supernatant was measured by ELISA at 6hpi. ns – not significant, * $p < 0.05$, ** $p < 0.01$, *** $p < 0.001$, **** $p < 0.0001$ by Tukey's (B) or Dunnett's (C) multiple comparisons test. Error bars represent the standard deviation of triplicate wells from one experiment. Data shown are representative of at least three independent experiments.

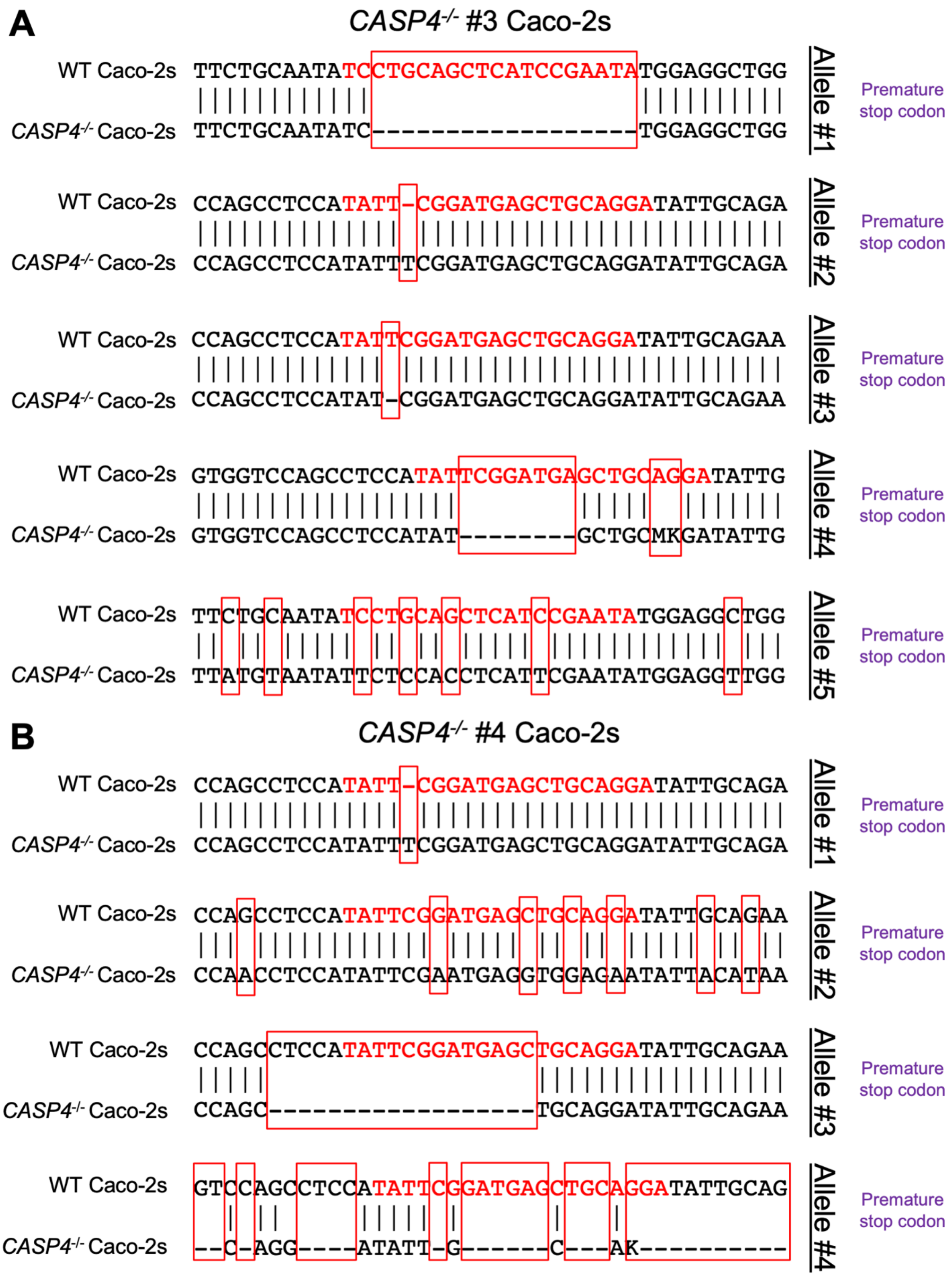
A
CASP4-/- #3 Caco-2s
WT Caco-2s TTCTGCAATATCCTGCAGCTCATCCGAATATGGAGGCTGG
CASP4-/- Caco-2s TTCTGCAATATC------------------TGGAGGCTGG
Allele #1
Premature stop codon
WT Caco-2s CCAGCCTCCATATT-CGGATGAGCTGCAGGATATTGCAGA
CASP4-/- Caco-2s CCAGCCTCCATATTTCGGATGAGCTGCAGGATATTGCAGA
Allele #2
Premature stop codon
WT Caco-2s CCAGCCTCCATATTCGGATGAGCTGCAGGATATTGCAGAA
CASP4-/- Caco-2s CCAGCCTCCATAT-CGGATGAGCTGCAGGATATTGCAGAA
Allele #3
Premature stop codon
WT Caco-2s GTGGTCCAGCCTCCATATTCGGATGAGCTGCAGGATATTG
CASP4-/- Caco-2s GTGGTCCAGCCTCCATAT--------GCTGCMKGATATTG
Allele #4
Premature stop codon
WT Caco-2s TTCTGCAATATCCTGCAGCTCATCCGAATATGGAGGCTGG
CASP4-/- Caco-2s TTATGTAATATTCTCCACCTCATTCGAATATGGAGGTTGG
Allele #5
Premature stop codon
B
CASP4-/- #4 Caco-2s
WT Caco-2s CCAGCCTCCATATT-CGGATGAGCTGCAGGATATTGCAGA
CASP4-/- Caco-2s CCAGCCTCCATATTTCGGATGAGCTGCAGGATATTGCAGA
Allele #1
Premature stop codon
WT Caco-2s CCAGCCTCCATATTCGGATGAGCTGCAGGATATTGCAGAA
CASP4-/- Caco-2s CCAACCTCCATATTCGAATGAGGTGGAGAATATTACATAA
Allele #2
Premature stop codon
WT Caco-2s CCAGCCTCCATATTCGGATGAGCTGCAGGATATTGCAGAA
CASP4-/- Caco-2s CCAGC------------------TGCAGGATATTGCAGAA
Allele #3
Premature stop codon
WT Caco-2s GTCCAGCCTCCATATTCGGATGAGCTGCAGGATATTGCAG
CASP4-/- Caco-2s --C-AGG----ATATT-G------C---AK----------
Allele #4
Premature stop codon

**Figure S10 (related to Figure 7): Validation of *CASP4* mutant Caco-2 cells generated with CRISPR/Cas9 genome editing.** Sequence alignments of WT Caco-2, *CASP4*$^{-/-}$ #3, and *CASP4*$^{-/-}$ #4 Caco-2 are shown for multiple alleles. Red text represents the target guide RNA sequence and red boxes represent the mutated region. Purple text represents the predicted impact of the mutation on the amino acid sequence.

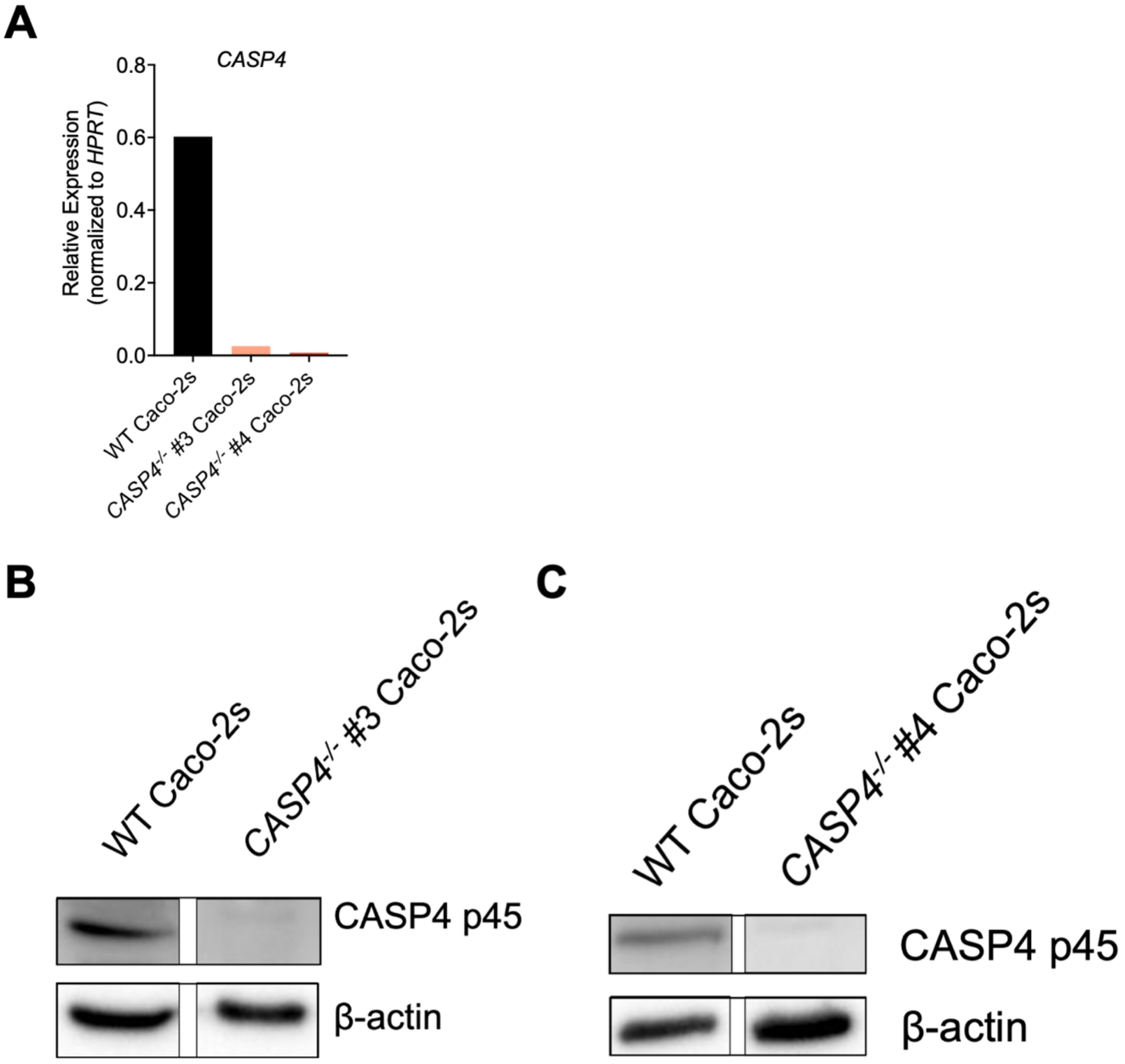


**Figure S11 (related to Figure 7): Validation of *CASP4* mutant Caco-2 cells generated with CRISPR/Cas9 genome editing.** (A) Relative mRNA expression of

*CASP4* compared to the housekeeping control *HPRT* in WT Caco-2s, *CASP4*$^{-/-}$ #3, and *CASP4*$^{-/-}$ #4 Caco-2s, as measured by qRT-PCR. (B, C) WT Caco-2s, *CASP4*$^{-/-}$ #3, and *CASP4*$^{-/-}$ #4 Caco-2s lysates were immunoblotted for CASP4 and β-actin.

Supplemental Figure Legends

**Figure S1 (related to Figure 1): *Salmonella* infection induces IL-8 release in human intestinal epithelial cells.** (A) Caco-2 cells or (B) T84 cells were infected with PBS (Mock), WT *S*. Typhimurium, or Δ*sipB S*. Typhimurium. Release of IL-8 into the supernatant was measured by ELISA at 6hpi. (C) Caco-2 cells were treated with 30 μM disulfiram or DMSO as a vehicle control 1 hour prior to infection. Cells were then infected with PBS (Mock), WT *S.* Typhimurium*,* or Δ*sipB S*. Typhimurium. Release of IL-8 into the supernatant was measured by ELISA at 6hpi. ns – not significant, * $p < 0.05$, ** $p < 0.01$, **** $p < 0.0001$ by Dunnett's multiple comparisons test (A), or by unpaired t-test (B), or by Šídák's multiple comparisons (C). Error bars represent the standard deviation of triplicate wells from one experiment. Data shown are representative of at least three independent experiments.

**Figure S2 (related to Figure 2): Bacterial T3SS ligands delivered using *Listeria* do not activate the inflammasome in human intestinal epithelial cells.** WT Caco-2 cells (A), T84 cells (B), polarized Caco-2 cells (C) or C2Bbe1 cells (D) were primed for 3h with 400 ng/ml of Pam3CSK4 and infected with PBS (Mock), WT *L. monocytogenes* (WT Lm), or *L. monocytogenes* expressing *S.* Typhimurium SPI-1 inner rod (Lm + PrgJ), or SPI-1 needle (Lm + PrgI) at the following MOIs: Caco-2 = 5; T84 = 25; polarized Caco-2 = 100; C2Bbe1 = as indicated in graph. Release of IL-18 into the supernatant was measured by ELISA at 16hpi. ns – not significant, ** $p < 0.01$, **** $p < 0.0001$ by Dunnett's multiple comparisons test. L.O.D indicates the limit of detection of the assay.

Error bars represent the standard deviation of triplicate wells from one experiment. (B) and (D) were conducted in duplicate wells.

**Figure S3 (related to Figure 2): Bacterial T3SS ligands delivered using the anthrax toxin system do not activate the inflammasome in human intestinal epithelial cells.** WT Caco-2 cells (A), polarized Caco-2 cells (B), C2Bbe1 cells (C), or T84 cells (D) were primed for 3h with 400 ng/ml of Pam3CSK4. Cells were then treated with PBS (Mock), PA alone, LFnPrgJ alone, or PA+LFnPrgJ (PrgJTox) for 16 hours. Release of IL-18 into the supernatant was measured by ELISA. For polarized Caco-2s (B), supernatants were collected from both apical and basolateral compartments. L.O.D indicates the limit of detection of the assay. Error bars represent the standard deviation of triplicate wells from one experiment. (A, C, D) were conducted in duplicate wells. Data shown are from one independent experiment (A, D) or are representative of two independent experiments (B, C). ns – not significant, ** $p < 0.01$ by Dunnett's multiple comparisons test.

**Figure S4 (related to Figure 4): Validation of *NAIP* mutant Caco-2 cells generated with CRISPR/Cas9 genome editing.** Sequence alignments of WT Caco-2, *NAIP*$^{-/-}$ #7 Caco-2, and *NAIP*$^{-/-}$ #8 Caco-2 are shown for multiple alleles. Red text represents the target guide RNA sequence and red boxes represent the mutated region. Purple text represents the predicted impact of the mutation on the amino acid sequence.

**Figure S5 (related to Figure 5): Validation of *PYCARD* mutant Caco-2 cells generated with CRISPR/Cas9 genome editing.** Sequence alignments of WT Caco-2, *PYCARD*$^{-/-}$ #4, and *PYCARD*$^{-/-}$ #6 Caco-2 are shown for multiple alleles. Red text represents the target guide RNA sequence and red boxes represent the mutated region. Purple text represents the predicted impact of the mutation on the amino acid sequence.

**Figure S6 (related to Figure 5): Expression of ASC in *PYCARD* mutant Caco-2 cells generated with CRISPR/Cas9 genome editing.** (A) Relative mRNA expression of *PYCARD* compared to the housekeeping control *HPRT* as measured by qRT-PCR in WT and *PYCARD*$^{-/-}$ #4 Caco-2s and *PYCARD*$^{-/-}$ #6 Caco-2s. (B) WT THP-1s, WT Caco-2s, *PYCARD*$^{-/-}$ #4 Caco-2s, and *PYCARD*$^{-/-}$ #6 Caco-2s lysates were immunoblotted for ASC and β-actin.

**Figure S7 (related to Figure 6): Validation of *CASP1* mutant Caco-2 Clone 7 generated with CRISPR/Cas9 genome editing.** (A) Sequence alignments of WT Caco-2s, and *CASP1*$^{-/-}$ #7 Caco-2s are shown for multiple alleles. Red text represents the target guide RNA sequence and red boxes represent the mutated region. Purple text represents the predicted impact of the mutation on the amino acid sequence. (B) Relative mRNA expression of *CASP1* compared to the housekeeping control *HPRT* in WT Caco-2s and *CASP1*$^{-/-}$ #7 Caco-2s, as measured by qRT-PCR. (C) WT THP-1s, WT Caco-2s, *CASP1-/-* #7 Caco-2 lysates were immunoblotted for ASC and β-actin.

**Figure S8 (related to Figure 6): Validation of *CASP1* mutant Caco-2 Clone 12 generated with CRISPR/Cas9 genome editing.** Sequence alignments of WT Caco-2, and *CASP1*$^{-/-}$ #12 Caco-2 are shown for multiple alleles. Red text represents the target guide RNA sequence and red boxes represent the mutated region. Purple text represents the predicted impact of the mutation on the amino acid sequence.

**Figure S9 (related to Figure 7): *Salmonella* infection induces CASP4/5-dependent inflammasome activation in human intestinal epithelial cells.** (A, B) WT Caco-2 cells were treated with siRNA targeting *CASP4, CASP5,* or a control scrambled siRNA for 72h. Cells were primed with 400 ng/ml of Pam3CSK4 for 3hrs. Cells were then infected with PBS (Mock), WT *S*. Typhimurium*,* or Δ*sipB S*. Typhimurium for 6hrs. (A) Knockdown efficiency was measured by qRT-PCR and normalized to housekeeping gene *HPRT*, and calculated relative to control-siRNA-treated cells. (B) Release of IL-8 into the supernatant was measured by ELISA at 6hpi. (C) WT or two independent clones of *CASP4*$^{-/-}$ Caco-2 cells were infected with PBS (Mock), WT *S*. Typhimurium, or Δ*sipB S*. Typhimurium for 6hrs. Release of IL-8 into the supernatant was measured by ELISA at 6hpi. ns – not significant, * $p < 0.05$, ** $p < 0.01$, *** $p < 0.001$, **** $p < 0.0001$ by Tukey's (B) or Dunnett's (C) multiple comparisons test. Error bars represent the standard deviation of triplicate wells from one experiment. Data shown are representative of at least three independent experiments.

**Figure S10 (related to Figure 7): Validation of *CASP4* mutant Caco-2 cells generated with CRISPR/Cas9 genome editing.** Sequence alignments of WT Caco-2,

*CASP4$^{-/-}$* #3, and *CASP4$^{-/-}$* #4 Caco-2 are shown for multiple alleles. Red text represents the target guide RNA sequence and red boxes represent the mutated region. Purple text represents the predicted impact of the mutation on the amino acid sequence.

**Figure S11 (related to Figure 7): Validation of *CASP4* mutant Caco-2 cells generated with CRISPR/Cas9 genome editing.** (A) Relative mRNA expression of *CASP4* compared to the housekeeping control *HPRT* in WT Caco-2s, *CASP4$^{-/-}$* #3, and *CASP4$^{-/-}$* #4 Caco-2s, as measured by qRT-PCR. (B, C) WT Caco-2s, *CASP4$^{-/-}$* #3, and *CASP4$^{-/-}$* #4 Caco-2s lysates were immunoblotted for CASP4 and β-actin.
